# Spatial and Molecular Progression of Neural Progenitor Cells in the Developing Human Dentate Gyrus

**DOI:** 10.64898/2026.06.06.730648

**Authors:** Oier Pastor-Alonso, Matthew G. Heffel, Mohammad S. Baig, Jasmine Harris, Susana González-Granero, Siqian Li, Sol Beccari, Julia Chu, Hannah Lambing, I-Ling Lu, Matthew Varughese, Andrew L. Cheng, Jessica Le, Mohini Bhade, JaeYeon Kim, Arantxa Cebrian-Silla, Isabel Torres-Cuevas, Kurtis I. Auguste, Eric Huang, Arturo Alvarez-Buylla, Jesse Gomez, José Manuel Garcia-Verdugo, Chongyuan Luo, Mercedes F. Paredes

**Affiliations:** Department of Neurology, University of California San Francisco, San Francisco, CA 94143, USA; Weill Institute for Neurosciences, University of California San Francisco, San Francisco, CA 94143, USA; Department of Human Genetics, University of California Los Angeles, Los Angeles, CA 90095, USA; Bioinformatics Interdepartmental Program, University of California Los Angeles, Los Angeles, CA 90095, USA; Genomic Analysis Laboratory, The Salk Institute for Biological Studies, La Jolla, CA 92037, USA; Laboratory of Comparative Neurobiology, Cavanilles Institute of Biodiversity and Evolutionary Biology, University of Valencia and CIBERNED-ISCIII, Valencia, Spain; Preclinical Therapeutic Testing Core in the Brain Tumor Center, University of California San Francisco, San Francisco, CA 94143, USA; The Eli and Edythe Broad Center of Regeneration Medicine and Stem Cell Research, University of California, San Francisco, San Francisco, CA 94143, USA; Department of Physiology, Faculty of Medicine, University of Valencia, Valencia, Spain; Neonatal Research Group, Health Research Institute La Fe (IIS La Fe), Av. Fernando Abril Martorell 106, Valencia, Spain; Department of Neurological Surgery, University of California San Francisco (UCSF), San Francisco, CA, United States; Department of Pathology, University of California San Francisco, San Francisco, CA, USA; Pathology Service, San Francisco VA Health Care System, San Francisco, CA, USA; Department of Pathology & Immunology, Washington University School of Medicine, St. Louis, MO, USA; Princeton Neuroscience Institute, Princeton University, Princeton, NJ, USA; Developmental Stem Cell Biology, University of California San Francisco, San Francisco, CA 94143, USA

**Author notes:** C.L. and M.F.P. are Co-corresponding authors. O.P.A and M.G.H. are Co-first authors.

## Abstract

The large diversity of neuronal and glial cell types in the human brain is underpinned by foundational cell populations known as neural progenitor cells (NPCs). The dentate gyrus (DG) of the hippocampus, a key structure in learning and memory, maintains a tightly organized NPC population into adulthood across many mammalian species. However, the emergence, organization and persistence of NPCs in the human hippocampus remain poorly characterized. Reports of NPCs in the juvenile, adult, and aged periods have been variable, reflecting differences in identification criteria and highlighting the need for a unified framework across development. In this study, we provide a spatial and molecular map of the developmental trajectory of NPCs in the human DG, combining multimodal transcriptomic analysis within a neuroanatomical context. At mid-gestation, we observed changes in the structural and cellular arrangement of the hippocampus, coinciding with the emergence of a multicellular NPC layer within the DG, herein named the granular-hilar progenitor zone (GHPZ). Neurogenic transcriptomic signatures in the GHPZ were diminished by early infancy, coinciding with a reduction in NPC number as they progressed toward an astrocytic program. At childhood, the GHPZ dissolved with only sparse radial NPCs remaining in the DG. Lastly, we validated WNT signaling pathway-associated genes as NPC identity markers in the developing human DG, observing a decline in their expression after infancy. Our study defines the steep decline of NPCs from gestation to the postnatal period, identifies their progression to an astrocytic nature, and sets the molecular blueprint for NPC identification in the human DG.

**Highlights:**

- Multimodal mapping of neural progenitor cells from gestational to postnatal stages in the human hippocampus
- Formation of the granular-hilar progenitor zone within the dentate gyrus at mid-gestation
- Neurogenic potential declines sharply from the prenatal period to childhood, with radial glia cells progressively acquiring astrocytic features
- Developmental modulation of the WNT signaling pathway accompanies radial glia cell transitions

## Introduction

During development, the complex neuronal and glial mosaic that underpins brain functionality is generated from foundational cell populations known as neural progenitor cells (NPCs). Among these, radial glial cells (RGCs) function as the primary progenitors, directly giving rise to neurons and glia while also generating secondary progenitor populations, known as intermediate progenitor cells (INPs)^1–4^. In most brain regions, NPCs are transient populations that disappear by birth. However, the mammalian dentate gyrus (DG), a key structure to regulate cortical inputs into the hippocampus^5,6^, retains NPCs through adulthood in rodents and non-human primates^7–9^. This capacity to generate neurons beyond the developmental window has attracted significant interest in the last decades, owing to its relevance as a plasticity mechanism and therapeutic potential. Investigating the early developmental dynamics of NPCs is central to understanding their function after birth.

Insights into NPC organization in the developing DG have largely come from rodent studies, which have defined the developmental sequence of DG in detail. During the embryonic period, the dentate ventricular zone (DVZ), a specialized portion of the hippocampal ventricular surface, harbors the earliest DG-bound NPCs. The DVZ progressively gives rise to a second NPC niche, the dentate migratory stream (DMS), a defined tangential corridor that directs migrating cells into the forming DG before and early after birth^10–12^. In parallel, the fimbria provides a gliogenic structural substrate that interfaces with the DMS and contributes to the DG cellular assembly^13,14^. By the first two weeks after birth, NPCs disappear from the DVZ and DMS and organize a final niche within the DG that is maintained long-term into adulthood, named the subgranular zone^10,15^. Together, these transient niches establish the canonical framework for DG development in rodents. Human DG development shows both conserved and divergent features relative to rodents. Different studies have reported a DVZ-DMS-DG migratory process comparable to the rodent system^16–18^, although questions remain on the temporal and structural basis of the different NPC niches. The presence of an NPC structure equivalent to the rodent subgranular zone, described in non-human primates^8,9,19^, has not been observed in humans, suggesting evolutionary divergence in the underlying developmental programs.

The presence of NPCs in the adult human DG has long been debated. Histological studies have yielded contradictory results^16,18,20–22^, partially driven by disparities in diverging criteria for molecular marker expression, morphological identification and spatial location of NPCs. Transcriptomic studies have described NPC signatures in gestation and infancy^23,24^, but their identification in adulthood has proven more challenging. Machine-learning approaches have been used to identify NPC signatures in the adult, aged and pathological DG^24,25^. However, the results differ in NPC numbers and in the gene signatures reported. Moreover, studies identifying NPCs transcriptomically have relied on canonical molecular markers without spatial or morphological validation. Together, these observations underscore the need for an integrated framework to resolve the defining features and developmental progression of NPCs in the human DG.

To address these gaps, we generated a multimodal atlas of the developing human DG, integrating high-resolution imaging, spatial transcriptomics and single-nucleus RNA-sequencing across gestation, infancy, and childhood. By aligning molecular identity, spatial localization, and morphology, we characterized NPC populations that contribute to DG formation and define their lineage transitions during the pre- and postnatal periods. Our work establishes the foundational principles of human DG development, resolving the organization and temporal progression of NPCs.

## Results

### Cytoarchitectural organization of the neurogenic niches

Through evolution, the hippocampus undergoes a dramatic reorientation from a vertical configuration in rodents to a horizontal configuration in humans^26^. Given the relationship between developmental programs and phylogenetic differences across species^27^, we measured positional and volumetric changes of the human developing hippocampus to provide a gross anatomical context for NPC progression. We segmented the hippocampus in 3D space using an in vivo magnetic resonance imaging dataset that spanned the gestational time period of 21 to 38 gestational weeks (GWs)^28^. To provide a reference of adulthood, we employ the Montreal Neurological Institute adult brain template, an anatomical standard in neuroimaging produced from the volumetric averaging of 152 brains^29^ **(Figure 1A)**. First, we calculated the volume of the hippocampus across gestation, observing steady hippocampal growth, although outpaced by the expansion of the temporal cortex **(Figures 1B)**. Next, we measured the angle formed by the longest axis of the hippocampus and that of the anterior and posterior commissures, a normalized measurement of medial temporal lobe orientation. The results revealed early ventral rotation and medial–lateral repositioning of the hippocampus. Mid-gestation was the period of most rapid structural change, marked by a sharp decline in the hippocampal angle relative to the anterior–posterior commissure axis, followed by a period of more gradual angular change. This pattern of angular decay was well-described by an exponential decay model (R^2^ = 0.93) **(Figure 1C)**. To identify the transition point from this rapid to gradual angular development, we performed a two-lines test^30^, finding significant statistical evidence for an inflection point at 25 gestational weeks (slopes = −4.83 and 0.72, p<0.001) **(Figure 1C)**. These data suggest that mid-gestation represents a structural inflection point, during which the rapid cortical expansion unique to humans coincides with hippocampal rotation, providing a macroanatomical developmental context in which to investigate whether similarly unique routines occur at the cellular level.

**Figure 1.**
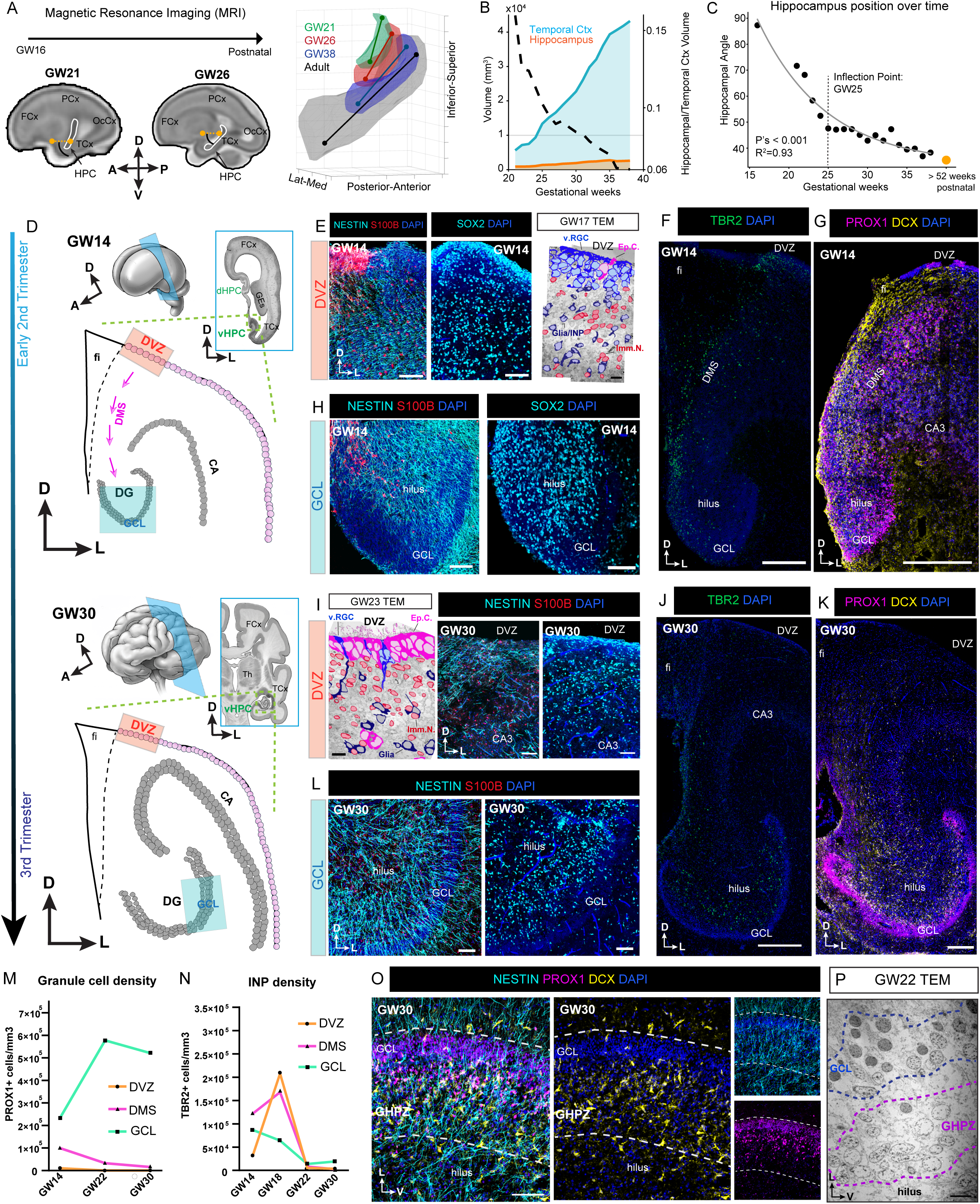
Anatomical and cytoarchitectural characterization of the human hippocampus from early- to mid- gestation. **(A)** Left: Representative MRI scans from GW21 and GW26 specimens with the HPC highlighted; the yellow line indicates the anterior commissure–posterior commissure plane used for hippocampal angle measurements. Right: Three-dimensional reconstructions of the hippocampus at GW21, GW26, GW38, and an adult reference from the MNI-152 average brain. **(B)** Volumetric changes in the HPC, frontal and temporal cortices over development. **(C)** Change in hippocampal angle relative to the anterior and posterior commissure (AC-PC) plane across development. Two lines test identified 25.5GW as an inflection point in angle change. **(D)** Schematics of the developing HPC at GW14 and GW30. Colors highlight the main neurogenic niches driving the formation of the HCX (DVZ, DMS and DG). **(E)** Left: immunofluorescence for NESTIN and S100B in the DVZ at GW14. Middle: Immunofluorescence for SOX2 in the ventral DVZ at GW14. Right: Outlines of cell types identified on ultrathin sections by TEM in the DVZ at GW17. Ventricular RGCs (blue), a heterogeneous population of glial cells and INPs (dark blue), ependymal cells (magenta) and immature neurons (red) classified based on their ultrastructural characteristics within the DVZ and along their migratory route toward the DMS. **(F-G)** Immunofluorescence for TBR2, PROX1 and DCX on the GW14 hippocampus. **(H)** Immunofluorescence for NESTIN and S100B (left) and SOX2 (right) in the DG at GW14. **(I)** Left: Outlines of cell types identified on ultrathin sections by TEM in the DVZ at GW23. Ventricular RGCs (blue), glial cells (dark blue), ependymal cells (magenta) and immature neurons (red) classified based on their ultrastructural characteristics within the remaining DVZ. Middle: Immunofluorescence for NESTIN and S100B in the DVZ at GW30. Right: Immunofluorescence for SOX2 in the DVZ at GW30 **(J-K)** Immunofluorescence for TBR2 (J) and PROX1 and DCX (K) on the GW30 hippocampus. **(L)** Immunofluorescence for NESTIN and S100B (left) and SOX2 (right) in the DG at GW30. **(M-N)** Quantification of TBR2^+^ and PROX1^+^ cell densities in the DVZ, DMS, and DG from GW14 to GW30. Data are presented as the mean of individual measurements across 2 to 4 independent sections per case. Each age is represented by a n=1 case. **(O)** Immunofluorescence for NESTIN, PROX1 and DCX in the GHPZ and GCL at GW22 and GW30. (P) TEM images of the GHPZ and GCL area at GW22. <u>Abbreviations:</u> CA, cornu ammonis; D, dorsal; DG, dentate gyrus; DMS, dentate migratory stream; DVZ, dentate ventricular zone; Fi, fimbria; FCx, frontal cortex; GCL, granule cell layer; GE, ganglionic eminence; GHPZ, granular-hilar progenitor zone; GW, gestational week; HPC, hippocampus; L, lateral; OcCx, occipital cortex; PCx, parietal cortex; TCx, temporal cortex; V, ventral; VZ, ventricular zone. <u>Scale bars:</u> (E, H, I, L, O) 100 μm; (F, G, J, K) 300 μm; (E, I, TEM) 10 μm.

To define the cytoarchitectural basis underlying the observed anatomical changes at mid-gestation, we examined coronal sections at four different ages (GW14, GW18, GW22 and GW30) and characterized the neurogenic niches within the main regions driving DG formation (the DVZ, DMS and DG itself) **(Figure 1D)**. Using immunohistochemistry and canonical markers, we mapped the full neurogenic cascade from RGCs to INPs, immature migratory neurons, granule neurons, and astrocytes^15^. From GW14 (both in the dorsal and ventral hippocampi present at this age) to GW18, RGCs, defined by NESTIN expression and radial morphology^31^, occupied the entire hippocampal formation. Co-expression of NESTIN and S100B, a marker for cells in the astrocytic, oligodendrocytic and ependymal lineage^32,33^, was only observed in the fimbria, highlighting the glial nature of this area **(Figure S1A)**. Immediately adjacent to the fimbria, the DVZ was identified as a dense cellular clustering with numerous NESTIN^+^ processes that extended toward the DMS, and the additional RGC marker SRY-box transcription factor 2 (SOX2)^34^ corroborated the continuous presence of RGCs throughout the DVZ and DMS **(Figures 1E and S1B)**. Transmission electron microscopy (TEM) analysis of the DVZ at GW17 revealed RGCs attached to the ventricular surface, a heterogeneous population of glial cells and INPs detached from the ventricle, and abundant migratory cells, identified by their elongated morphology and a leading process oriented toward the DMS **(Figures 1E and S1B)**. The neurogenic activity of the DVZ-DMS at this age was confirmed by a dense cohort of t-box brain transcription factor 2 (TBR2)^+^ cells, a canonical marker for INPs^35^, distributed in a continuous trajectory all along the DMS **(Figure 1F and S1C)**. Ki67^+^ proliferating cells and ultrastructurally-defined mitotic figures associated with migratory cells were similarly observed within this region **(Figure S1D)**, and the presence of doublecortin (DCX)^+^ migrating immature neurons^36^ and cells expressing prospero homeobox 1 (PROX1), associated with granule neuron lineage within the DG^37^, suggests granule neuron identity as the terminal fate of DVZ- and DMS-resident NPCs **(Figure 1G and S1E)**. NESTIN^+^ and SOX2^+^ RGCs connected the DMS to the DG via a continuous cellular chain that extended into the granule cell layer (GCL) **(Figures 1H and S1F)**. Notably, homeodomain-only protein homeobox (HOPX), reported as a DG-specific RGC marker in the mouse^12^, was not expressed in RGCs in the developing human DG. Instead, HOPX was co-expressed with the astrocyte marker glial fibrillary acidic protein (GFAP) in cells restricted to the fimbria **(Figure S1G)**. Altogether, our data revealed the cytoarchitectural framework for cellular migration into the developing DG.

By mid-gestation (GW22 to GW30), while S100B-expressing cells remained limited to the fimbria, NESTIN expression was reduced to focal points of the hippocampal formation **(Figure S1H)**. Ultrastructural analysis of the DVZ confirmed that at GW17, the majority of cells lining the ventricular epithelium were RGCs, characterized by the presence of a primary cilium and a long radial expansion with intermediate filaments^38^. By GW22 this composition had shifted dramatically, with most cells differentiated into multiciliated ependymal cells^39–41^ **(Figure 1I and S1I; Table S2)**. Glial cells lacking ventricular contact were also observed, along with a small number of displaced ependymal cells, and chains of migratory neurons now oriented parallel to the ventricular wall **(Figure 1I and S1J)**. Through immunofluorescence, NESTIN^+^ and SOX2^+^ cells were observed forming a scaffold parallel to the ventricular surface, including at the remnant DVZ **(Figures 1I and S1K)**, and co-expression of DCX with the inhibitory neuron marker glutamate decarboxylase 1 (GAD1) revealed the inhibitory identity of the migratory neurons persisting in this region **(Figure S1L)**. Together, these features are consistent with a transition toward a more mature, non-neurogenic state of the DVZ. This maturation was confirmed by the absence of TBR2^+^, Ki67^+^ and PROX1^+^ cells from the DVZ and DMS by GW22-GW30 **(Figures 1J, 1K, S1M and S1N)**. In contrast, TBR2^+,^ Ki67^+^ and DCX^+^ cells, together with NESTIN^+^ and SOX2^+^ RGCs, organized in a diffuse layer between the hilus and GCL **(Figures 1J, 1K, S1L, S1M and S1N)**. The quantifications of TBR2^+^ and PROX1^+^ cell densities at the DVZ, DMS and GCL validated a progressive reduction of both TBR2^+^ and PROX1^+^ cells in the DVZ and DMS from GW14 to GW30 **(Figure 1M and 1N; Table S2)**. HOPX expression was widespread across the hippocampus, often within GFAP^+^ cells with astrocytic morphology, consistent with an astroglial identity **(Figure S1O)**. These results highlight mid-gestation as an inflection point for NPC reorganization, when DG-related neurogenesis transitions from the ventricular wall into a stratified cellular organization within the DG.

We further characterized the emergence of the NPC niche at the interface between the hilar region and the GCL. At GW22, we found abundant NESTIN^+^ RGCs extending their radial processes from the hilus toward the molecular layer, crossing the GCL. These RGCs were accompanied by DCX^+^ migrating neurons, and PROX1^+^ cells were gradually enriched at the hilus-GCL margin, reflecting the later stages of granule neuron differentiation. By GW30, this niche was further consolidated, with tight clusters of NESTIN^+^ RGCs intermingled with DCX^+^ immature neurons underneath a PROX1^+^ GCL and above a sparsely populated hilar region **(Figure 1O)**. Ultrastructural analysis of the hilus-GCL margin showed cells with sparse cytoplasm and few organelles situated beneath the mature GCL, consistent with an NPC-rich zone and confirming the cytoarchitectural organization of an emerging neurogenic niche within the DG **(Figure 1P)**. These results establish that a neurogenic structure forms between the GCL and the hilus of the DG at mid-gestation, herein termed the granular-hilar progenitor zone (GHPZ). This diffuse structure differed from the classic mammalian SGZ, in which stem cells are arranged in a thin, single-cell layer beneath the GCL^7^.

### Molecular profiling of the neurogenic niches

To capture the transcriptional programs that underlie the spatiotemporal shifts in NPCs, we performed spatially resolved bulk transcriptomics (GeoMx digital spatial profiling)^42^ in the human hippocampus at GW14, GW22-23, and 4 months **(Figure 2A)**. We manually segmented regions of interest (ROI) associated with the DG, including the DVZ, DMS, GHPZ, GCL, hilus and molecular layer. We also sampled CA-related regions at GW14, covering the ventricular wall through the CA neuronal layer, and the fimbria at all time points, to provide a broader spatial context for the interpretation of DG-related development. Additionally, two ROIs corresponding to the ganglionic eminence were captured in one GW23 sample. S100B and Vimentin were used as morphological markers for anatomical and cellular landmark identification **(Figures 2B and S2A; Table S3)**. This dataset provides a molecular map to study regional differences in the developing hippocampus.

**Figure 2.**
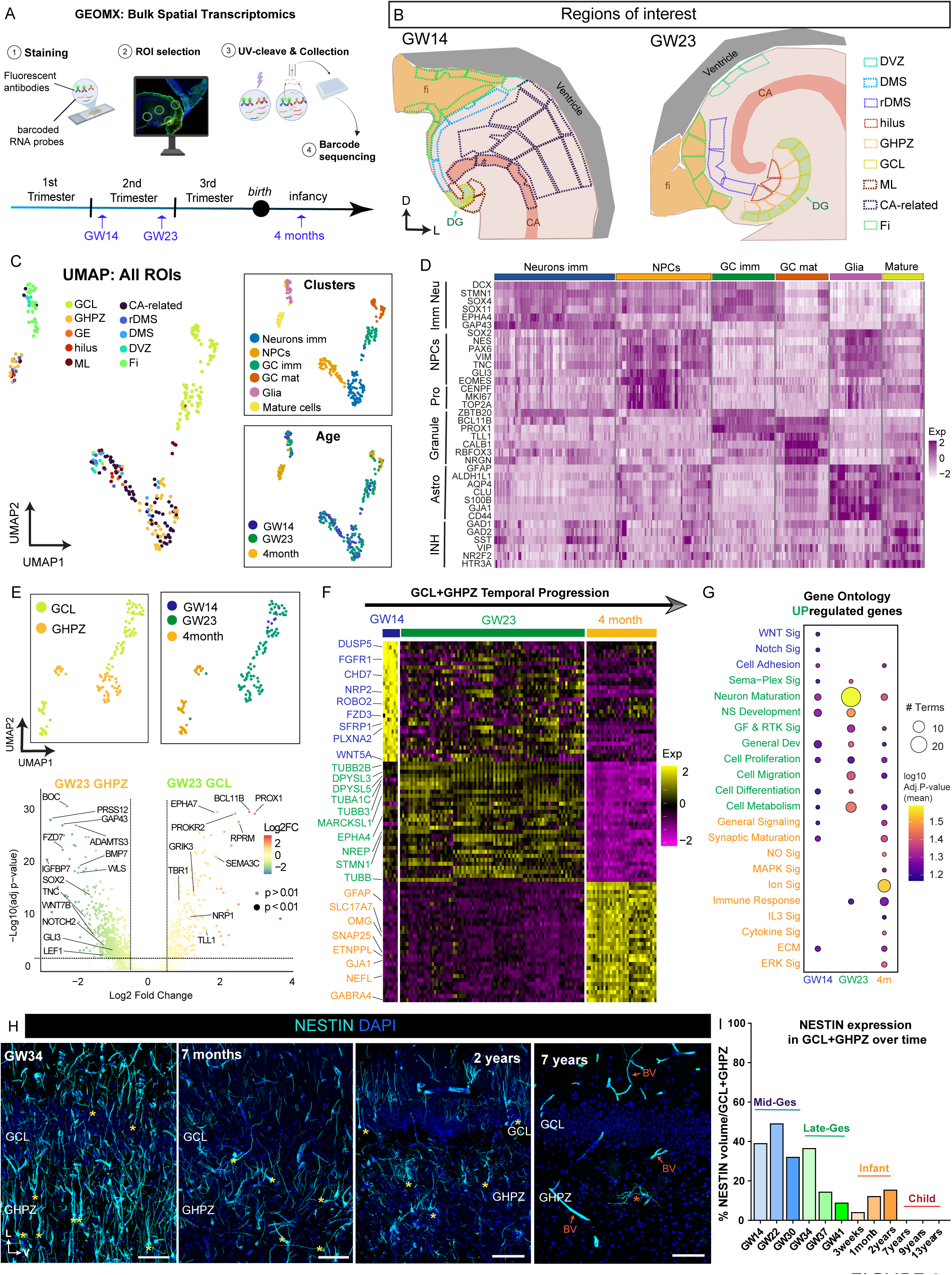
Spatial transcriptomic mapping of regional and neurogenic niche dynamics in the developing human hippocampus. **(A)** Schematic of GeoMx Digital Spatial Profiler workflow and human hippocampal cases analyzed (GW14, GW22-23 and 4 months). **(B)** Representative schematics of the regions of interest (ROIs) collected for GeoMx profiling at GW14 and GW23. Sampling at 4 months was equivalent to that at GW23. **(C)** Left: UMAP embedding of 280 ROIs annotated by anatomical region. Right: UMAPs annotated by unsupervised Leiden clustering (top) and age (bottom). **(D)** Heatmap of canonical markers for immature neurons, neural progenitor cells (NPCs), proliferating cells, granule neurons, astrocytes, and inhibitory neurons, illustrating the cellular composition of the clusters shown in (C). **(E)** Top: UMAP of 78 ROIs from the granule cell layer (GCL) and granular-hilar progenitor zone (GHPZ), annotated by region and developmental stage. Bottom: Volcano plot comparing GCL and GHPZ ROIs at GW22-23. Genes with adjusted p < 0.01 and |log₂ fold change| > 0.5 were considered differentially expressed. **(F)** Heatmap of the top upregulated genes in combined GCL and GHPZ ROIs at GW14, GW22-23, and 4 months. Selected neurodevelopmentally relevant genes are highlighted. **(G)** Gene Ontology biological process enrichment analysis of genes upregulated in combined GCL and GHPZ ROIs at GW14, GW22-23, and 4 months. **(H)** immunofluorescence for NESTIN in the GCL and GHPZ regions at GW34, 7 months, 2 years, and 7 years. Yellow asterisks highlight radial Nestin-expressing cells. Orange asterisk at 7 years highlight Nestin-expressing cells without radial morphology. Orange arrows point to Nestin-expressing blood vessels. **(I)** Quantification of the percentage of volume occupied by NESTIN immunoreactivity across 12 cases spanning GW14 to 13 years of age. The data is shown as the mean of at least three sections from distinct anterior–posterior levels per case. Each age is represented by a n=1 case. <u>Abbreviations</u>: BV, blood vessel; CA, cornu ammonis; D, dorsal; DG, dentate gyrus; DMS, dentate migratory stream; DVZ, dentate ventricular zone; Fi, fimbria; GCL, granule cell layer; GE, ganglionic eminence; GHPZ, granular-hilar progenitor zone; GW, gestational week; L, lateral; NPC, neural progenitor cell; ML, molecular layer; ROI, region of interest; UMAP, uniform manifold approximation and projection; V, ventral. <u>Scale bars:</u> (H) 100 µm.

We used this framework to interrogate molecular differences between regions. UMAP embedding and unbiased leiden clustering of 280 ROIs revealed six major clusters in our dataset **(Figure 2C)**. We used canonical marker gene expression, together with top-expressed genes, to annotate the distinct clusters. We found increased expression of the RGC markers NESTIN, TNC and PAX6, the INP marker EOMES, and the proliferation marker MKI67 in a cluster formed by DVZ and DMS regions from GW14 and GW22-23 samples and the CA ventricular zone at GW14 ROIs. Immature neuron gene signatures (DCX, STMN1, SOX4, SOX11) were enriched in the GHPZ ROIs at GW22-23, and CA-related ROIs at GW14, consistent with the presence of abundant immature neurons in these regions at these stages. Granule neuron signatures, with enriched expression of PROX1, were observed in two distinct clusters. One was formed by GW14 and GW22-23 GCL ROIs and showed expression of immature neuron genes such as DCX, STMN1, SEMA3C and EPHA7. The other was predominantly formed by 4M GCL ROIs and expressed more mature granule neuron markers, such as CALB1, RBFOX3, C1QL3 and CAMK2B, suggesting maturation changes in the GCL between mid-gestation and 4 months of age. Canonical signatures of diverse glial cell types, including P2RY12 (microglia), ALDH1L1 and S100B (astrocytes), OLIG2 (oligodendrocyte progenitor cells, OPCs), PECAM1 (endothelial cells) and FOXJ1 (ependymal cells), were observed in a clearly distinct cluster, formed by fimbria ROIs, further confirming the glial nature of this territory. Of note, inhibitory neuron genes, such as GAD1 and KCNS3, and glial gene expression were observed in a small cluster formed by ROIs from the 4 month sample, suggesting maturation changes in the early postnatal hippocampus **(Figures 2C, 2D and S2B)**. These results pinpoint the molecular tempo of differentiation across the principal territories of the developing hippocampus from mid-gestation to infancy and highlight early maturation in the GCL.

We next focused on the GHPZ, the neurogenic niche that persists within the DG after mid-gestation **(Figure 1)**. To identify key molecular drivers for the assembly and progression of this zone, we subset and re-analyzed 78 ROIs corresponding to the GHPZ and GCL **(Figure 2E)**. UMAP embedding showed clear distinctions between GW14 and GW22-23, and 4-month samples, as well as between GHPZ and GCL ROIs, supporting the distinct molecular identities of the two regions **(Figure 2E)**. Differential gene expression analysis between GHPZ and GCL at GW22-23 showed a more neuronal gene signature of the GCL (BCL11B, PROX1, SEMA3C or TLL1), whereas the GHPZ showed enrichment for RGC markers such as SOX2 and TNC and GLI3, and immature neuron markers such as GAP43, consistent with higher neurogenic potential and NPC presence in this region. Interestingly, WNT signaling pathway-related genes exhibited strong presence in the GHPZ, with FZD7, WLS, WNT7B and LEF1 enriched in this territory, hinting toward the relevance of this signaling pathway in the GHPZ **(Figure 2E)**. The temporal analysis of combined GHPZ and GCL regions demonstrated sharp molecular changes from GW22-23 to 4M. Differential gene expression between GW14, GW22-23 and 4 months showed progressive downregulation of neurogenesis-associated genes MARCKSL1, STMN1, EPHA4, NES, and upregulation of the astrocyte genes GFAP, S100B and ALDH1L1 **(Figure 2F, S2C, S2D)**, hinting toward loss of neurogenic potential in the DG by early postnatal stages. Gene ontology analyses of differentially expressed gene lists at each age revealed downregulation of terms associated to cell proliferation, differentiation, migration and metabolism at 4 months, coupled to upregulation of terms associated to cell structural maturation, cell functional maturation, synapse and ion signaling **(Figure 2G and S2E; Table S4)**. These results indicate a decrease in the neurogenic potential of the DG early after birth, in favor of cell maturation and circuit formation.

Next, we aimed to correlate the observed molecular progression in the GHPZ and GCL with changes in the cytoarchitecture of these territories. Ultrastructural analysis revealed consolidation of the GCL and absence of a laminar organization in the GHPZ at birth, accompanied by the appearance of synapses on somatic membranes, consistent with structural maturation **(Figure S2F)**. High-resolution confocal microscopy provided cellular-level insight into GHPZ maturation. At GW34, the GHPZ contained abundant NESTIN^+^ cells that extended a radial process toward the GCL, and NESTIN^+^ filaments were also observable in the GCL and molecular layer. During the infant period, the arrangement of NESTIN^+^ cells changed, with sparse clusters of NESTIN^+^ radial cells extending across the GHPZ and GCL. At 7 years of age, very sparse NESTIN^+^ radial cells were identified and were located across the DG, within the hilus, GCL, and molecular layer. NESTIN was expressed primarily by vascular populations **(Figures 2H)**, confirming a sharp NPC decline in the DG by childhood. Quantification of the volume occupied by NESTIN in 12 cases from GW14 to 13 years after birth confirmed a gradual decrease in immunoreactivity within the GHPZ and GCL over time **(Figure 2I; Table S2)**.

Notably, the medial portion of the DG showed a distinct spatial and temporal pattern. NESTIN⁺ cells with their filaments attached to the subpial zone connected with the medial GHPZ, forming a unique fibrous scaffold at mid- and late- gestation **(Figure S2I)**. Two years after birth, NESTIN^+^ cell clusters were observed in the medial arm of the DG in a more prominent fashion compared to the lateral region of the DG **(Figure S2I)**. However, these asymmetries were resolved by late childhood when the sparsity of NESTIN^+^ radial cells was homogeneous along the medial-lateral axis. Although NESTIN^+^ filamentous cells were observed in the fimbria, they lacked any spatial connection with the DG **(Figure S2I)**. These data highlight that molecular changes are coupled by cellular reorganization from mid-gestation to infancy, resulting in a progressive loss of NPCs and dissolution of the GHPZ by childhood.

### Transcriptional mapping of mid-gestation cell populations

To resolve how individual cell populations underpin the observed molecular changes in the DG, we performed highly multiplexed spatial transcriptomics imaging^43^ and profiled 52,115 single cells across multiple hippocampal regions at GW23 (**Figure 3A and S3A**). We identified 21 distinct clusters within the entire hippocampus, including RGCs, INPs, proliferating cells and major neuronal and glial populations. Within excitatory neuronal lineages, PROX1^+^ granule cells, two distinct SEMA3E^+^ CA3-hilus populations, two MPPED1^+^ CA1 populations, and two subicular populations, distinctively expressing BCL11B and FOXP1, and TSHZ2, respectively, were detected. We also identified MGE- and CGE-associated inhibitory neurons, with the latter further divided into VIP^+^ and VIP-populations. Cajal-Retzius cells expressing TP73 and RELN were located along the molecular layer of the DG, consistent with previous studies describing their role in guiding DG migration^44,45^. We identified oligodendrocyte precursor cells (OPCs) and glial progenitor cells (GPCs), defined by epidermal growth factor receptor (EGFR) enrichment^46,47^, widespread across the entire hippocampus. However, the fimbria showed a distinct population with astrocytic identity, supporting the presence of a distinct astroglial niche in this territory. Ependymal, endothelial, microglial and meningeal populations were also identified (**Figures 3B, 3C, S3B and S3C; Table S5**). These results expand the cellular characterization of the developing human hippocampus providing a spatially resolved molecular map of single cell populations in the mid-gestational hippocampus.

**Figure 3.**
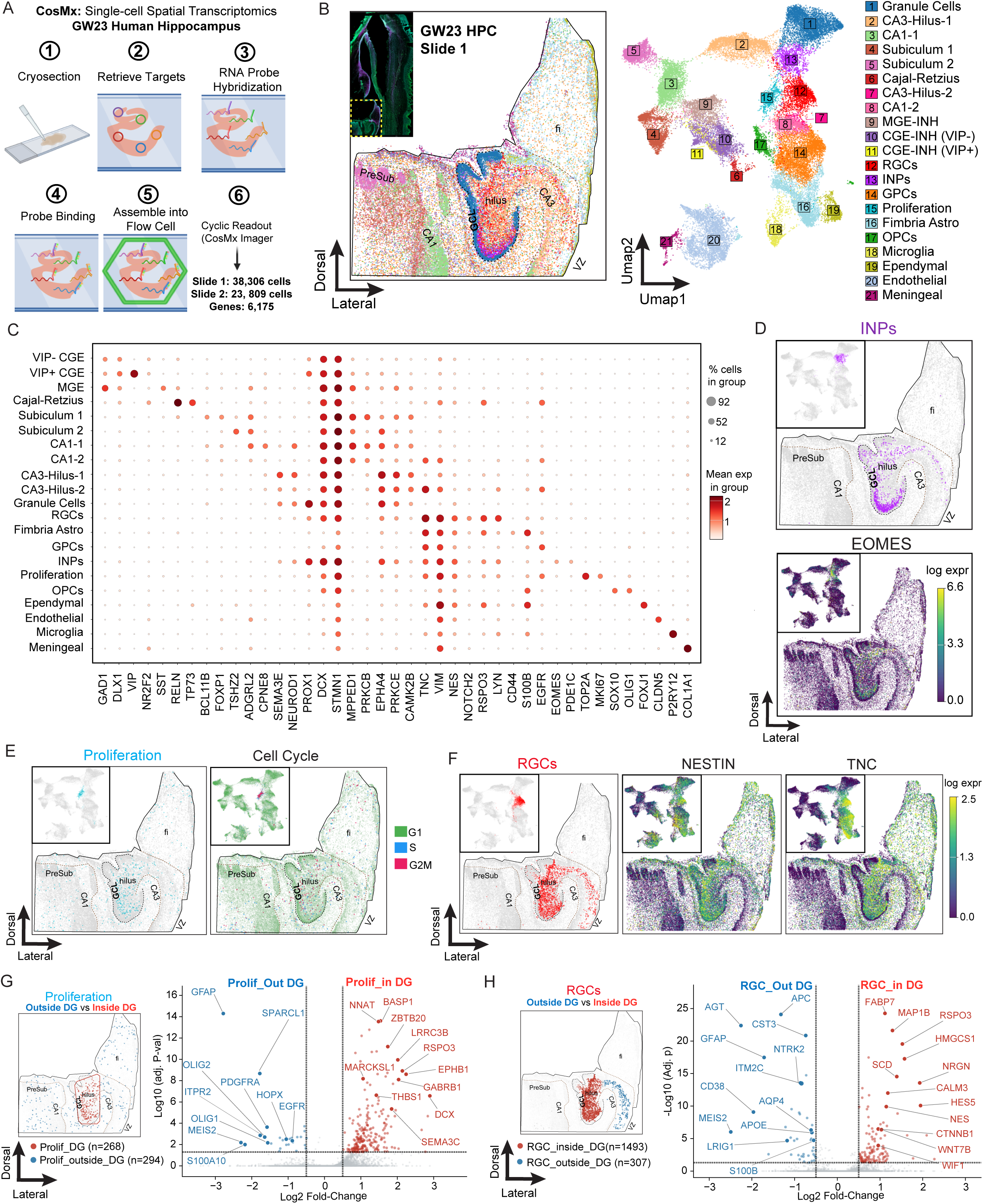
Single-cell spatial transcriptomic profiling of the human hippocampus at mid-gestation. **(A)** Schematic of CosMx Spatial Molecular Imager workflow applied to two sections from a GW23 human hippocampus. **(B)** Spatial mapping (left) and UMAP representation (right) of 21 individual cell types identified by unsupervised leiden clustering and manual annotation. **(C)** Dotplot highlighting the expression of canonical markers across the 21 clusters identified in B. **(D)** Spatial mapping of cells classified as INPs and the INP marker EOMES. **(E)** Left: Spatial mapping of cells classified as proliferating cells. Right: Cell cycle score analysis and spatial mapping of cells classified as G1, S or G2M phase cells. **(F)** Spatial mapping of cells classified as RGCs and the RGC markers NESTIN and TNC. **(G–H)** Left: Spatial mapping of proliferating cells (G) and radial glia-like cells (H) colored by location inside (red) or outside (blue) the DG. Right: Corresponding volcano plots comparing cells from inside versus outside the DG; genes with adjusted p < 0.05 and |log₂ fold change| > 0.5 were considered differentially expressed. <u>Abbreviations</u>: CA, cornu ammonis; CGE, caudal ganglionic eminence; fi, fimbria; GCL, granule cell layer; GE, ganglionic eminence; GHPZ, granular-hilar progenitor zone; GPC, glial progenitor cell; GW, gestational week; HPC, hippocampus; INP, intermediate neural progenitor; MGE, medial ganglionic eminence; RGC, radial glia cell; UMAP, uniform manifold approximation and projection; VZ, ventricular zone.

The spatial identification of NPCs and neurogenesis-related populations corroborated the cytoarchitectural rearrangement occurring at mid-gestation. INPs showed marked restriction into the GHPZ within the DG, confirmed by the expression pattern of EOMES **(Figures 3D and S3D)**. Proliferating cells were widespread across the hippocampus, with the GHPZ showing the highest proliferative potential. S- and G2M-phase cells identified through cell cycle score analysis corroborated this pattern **(Figures 3E and S3E)**. RGCs were also observed to be highly restricted to the DG, and canonical RGC markers (NESTIN, TNC) were primarily within the DG. However, these RGC markers showed strong expression in other glial populations, such as GPCs and fimbrial astrocytes (**Figure 3F and S3F**), highlighting the need for combined marker expression, spatial localization, and/or morphological criteria to identify RGCs. These results corroborate the formation of the GHPZ by mid-gestation with a detailed spatial and molecular identification of NPCs.

We further characterized NPCs from different spatial locations exploring molecular differences between populations within and outside the DG. First, we compared genetic signatures of proliferating cells. Cells outside the DG, occupying the territory between CA3, the ventricular wall and the fimbria, showed a marked oligodendrocytic identity with enriched expression of OLIG1, OLIG2 and PDGFRA. Astrocytic genes such as GFAP and SPARCL1 were also enriched in this group. In contrast, proliferating cells in the DG expressed neuronal genes such as DCX and SEMA3C **(Figures 3G and S3G)**, supporting the restriction of hippocampal neurogenic potential to the DG by mid-gestation. Consistently, RGCs outside the DG showed enriched expression of astrocytic genes, including GFAP, CST3, AQP4, APOE and S100B, whereas RGCs in the DG were enriched for FABP7 and NESTIN. Notably, WNT signaling-associated genes, such as R-spondin 3 (RSPO3), catenin beta 1 (CTNNB1), WNT7B and wnt-inhibitory factor 1 (WIF1), were also highly expressed in RGCs inside the DG compared to RGCs outside. These results suggest that RGCs outside the DG transition towards astrocytic lineages by mid-gestation, a change that parallels the loss of proliferative potential and is consistent with the relocation of NPCs to the GHPZ.

### Transcriptional trajectories of the cell populations

The above observations highlighted GW22-GW30 as a key time window for re-arrangement of DG-associated NPCs and the establishment of the GHPZ. To further profile the transcriptomic identities and trajectories governing the developmental progression of DG-associated cell populations, we performed single-nucleus RNA sequencing (snRNA-seq) on post-mortem human hippocampal samples spanning early-gestation (GW15, GW18, GW19), mid-gestation (GW21, GW23), infant (2 weeks, 4 months, 7 months), and adult (38 years) stages **(Figures 4A and S4A)**. Using manual cell annotation through unbiased leiden clustering and canonical marker expression, we identified NPC, macroglia, and major excitatory and inhibitory neurons superclasses. These superclasses were further subdivided into 19 cell classes, including RGCs, INPs and proliferating cells; astrocytes, oligodendrocytes, oligodendrocyte progenitor cells (OPCs) and glial progenitor cells (GPCs); CA pyramidal neurons, granule neurons, and immature neurons; and MGE-associated and CGE-derived inhibitory neurons, respectively. An additional superclass was termed “others”, including choroid plexus, endothelial cells, vascular leptomeningeal cells, microglia, ependymal cells, and less represented neuronal populations such as Cajal-Retzius neurons and thalamic neurons **(Figures 4A and S4B; Table S6)**. Canonical marker expression was used to delineate different cell types **(Figure 4B, 4C, S4C and S4D).** This dataset provides a comprehensive framework for the exploration of transcriptomic signatures in the developing DG.

**Figure 4.**
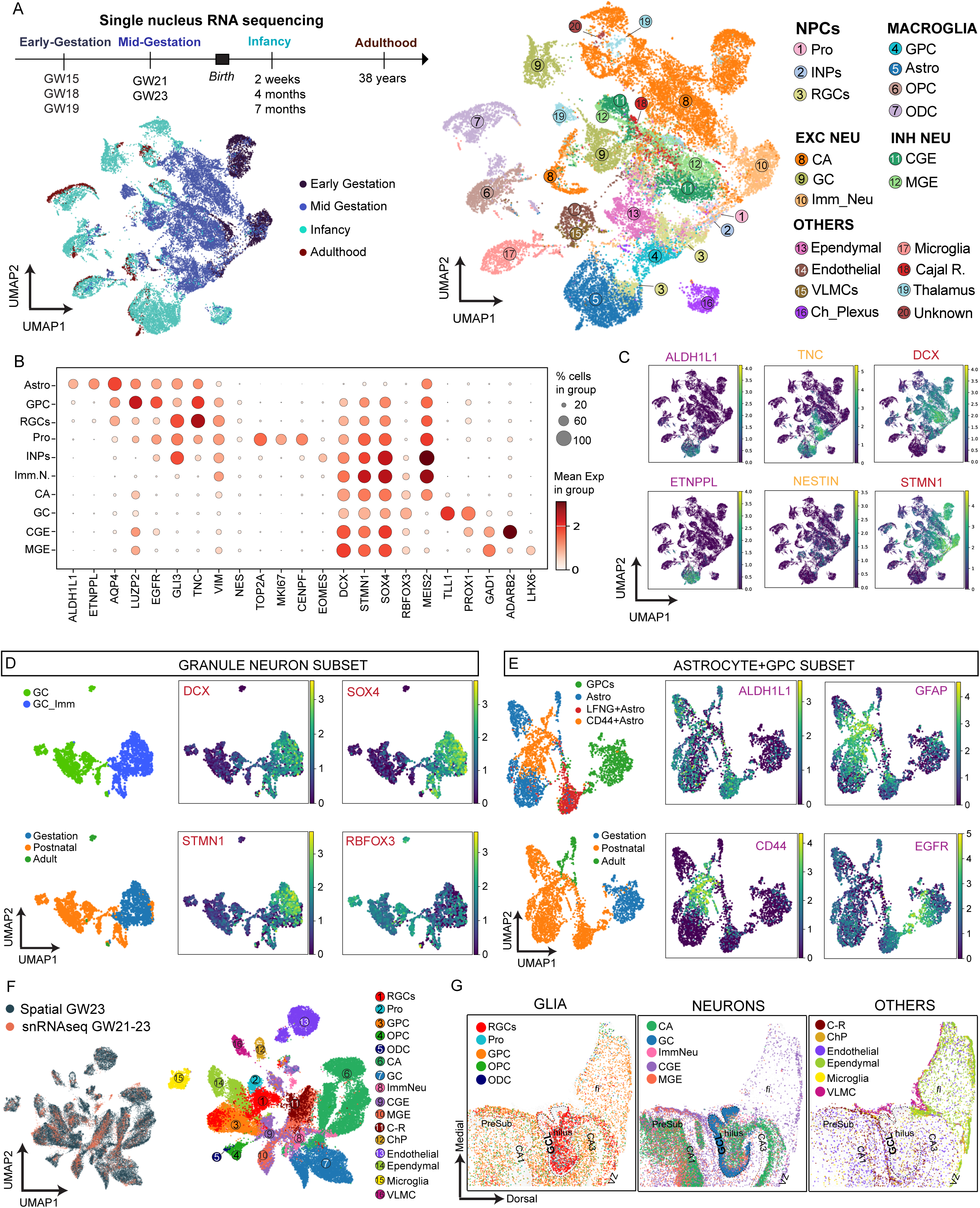
Single-nucleus transcriptomic profiling of the human developing hippocampus. **(A)** Timeline of single-nucleus RNA sequencing performed across nine samples from GW15 into adulthood (top left) and UMAP embeddings of 28,122 nuclei across nine hippocampi annotated by developmental stage (bottom left) and cell type classification (right). **(B)** Dotplot highlighting the expression of canonical markers across key cell types of interest identified in A. **(C)** UMAPs of canonical markers of astrocytes (ALDH1L1, ETNPPL), RGCs (TNC, NESTIN) and immature neurons (DCX, STMN1). **(D–E)** Left: UMAP embedding of nuclei classified as granule cells (D) and astrocytes and glial progenitor cells (GPCs) (E), annotated by cell subclass and developmental stage. Right: Expression of canonical markers defining each population, including immature (DCX, STMN1, SOX4) and mature (RBFOX3) neuronal markers in granule cells, and astrocyte (ALDH1L1, GFAP, CD44) and GPC (EGFR) markers in astrocyte/GPC populations. **(F)** Left: UMAP embedding of integrated spatial transcriptomic (GW23) and snRNA-seq datasets (GW21 and GW23). Right: Projection of snRNA-seq–defined cell types onto the integrated spatial–snRNA-seq embedding. **(G)** Spatial projection of snRNA-seq–defined cell types in the spatial context, with selected populations highlighted. <u>Abbreviations</u>: CA, cornu ammonis; CGE, caudal ganglionic eminence; fi, fimbria; GCL, granule cell layer; GHPZ, granular-hilar progenitor zone; GPC, glial progenitor cell; GW, gestational week; INP, intermediate neural progenitor; MGE, medial ganglionic eminence; RGC, radial glia cell; UMAP, uniform manifold approximation and projection; VZ, ventricular zone.

Next, we subset individual cell clusters and performed unsupervised subclustering to resolve subclasses within granule neurons, CA neurons, inhibitory neurons, and astrocytes. Across all neuronal populations, we identified subsets corresponding to immature (DCX, SOX4, STMN1) and mature (RBFOX3) neuronal states. However, these markers were not strictly restricted to discrete populations and instead showed varying levels of expression across both immature and mature subsets, reflecting the continuous maturation process that characterizes prenatal and early postnatal development **(Figures 4D, S4E and S4F)**. Granule neurons displayed the clearest maturation trajectory, with the immature subtype comprising predominantly gestational nuclei and a small subset of infant nuclei. The mature subtype included the remaining infant nuclei together with all adult nuclei. Within this mature population, adult nuclei clustered separately from infant nuclei in UMAP space **(Figure 4D)**. Immature CA neurons were detected throughout gestational, infant, and adult stages. Expression of immature markers (DCX, STMN1, SOX4) and the mature neuronal marker RBFOX3 overlapped across many nuclei, suggesting a gradual transition between maturation states rather than sharply defined populations **(Figure S4E)**. Similarly, inhibitory neurons exhibited a maturation gradient, although immature nuclei persisted into infancy and adulthood. Notably, expression of immature neuronal markers, particularly DCX, remained high in subsets of RBFOX3-high mature inhibitory neurons, suggesting the retention of immature molecular features for an extended developmental period within this lineage **(Figure S4F)**. We next subclustered astrocytes and GPCs together. GPCs were present at gestation and infancy, whereas astrocytes were only present in postnatal samples, delineating the astrocytic developmental trajectory. We observed three distinct astrocyte subclasses: LFNG-expressing astrocytes, only present at infant stages, CD44-expressing astrocytes and a canonical astrocyte group **(Figure 4E)**. These results shed light on different neuronal and astrocytic cell types present in the developing hippocampus, and define the developmental maturation trajectories from gestational to postnatal stages, highlighting a marked decline in immature neuronal signatures around the infant period.

To further validate the observed cell types, we integrated the GW23 and GW21 snRNA-seq samples with our GW23 spatial dataset and projected our snRNA-seq annotations onto the hippocampus **(Figure 4F and S4G)**. We observed OPCs, GPCs, microglia and endothelial cells scattered across the entire hippocampus, whereas ependymal cells were restricted to the fimbria and ventricular wall, and VLMCs to the subpial zone. Cajal-Retzius neurons showed major restriction to the molecular layer of the DG, and CGE- and MGE-derived inhibitory neurons were also widespread across the hippocampus, although with a slightly different spatial pattern. Granule neuron and CA neuron signatures mapped to their expected locations in the GCL and CA layer and hilus, respectively, and immature neurons showed a prominent mapping around CA, suggesting a predominant CA identity of this immature class. RGCs and proliferating cells showed significant presence within the hilus and GCL of the DG, supporting previous observations **(Figure 4G and S4H)**. These results provide spatial validation for our identified cell classes, including RGCs, and corroborate the relocation of NPCs toward the DG by mid-gestation.

### Transcriptional progression of NPCs

Collectively, our preceding observations underscored the formation of an NPC niche within the DG at mid-gestation, which gradually reduces its neurogenic capacity thereafter. To specifically study the transcriptomic progression of NPCs at the cellular level, we isolated RGCs from our snRNAseq dataset and interrogated their temporal transcriptomic progression **(Figure 5A)**. The subcluster analysis uncovered five transcriptionally distinct RGC subgroups—RGC1, RGC2, RGC2 astrocytic (RGC2A), RGC2 neuronal (RGC2N), and RGC3 **(Figure 5B)**. These subtypes mapped onto a defined developmental sequence: RGC1 predominated at early gestation; RGC2, RGC2A, and RGC2N appeared at mid-gestation, and RGC3 emerged after birth (**Figure 5C**). These results revealed distinct RGC transcriptomic identities from the early gestational period to infancy.

**Figure 5.**
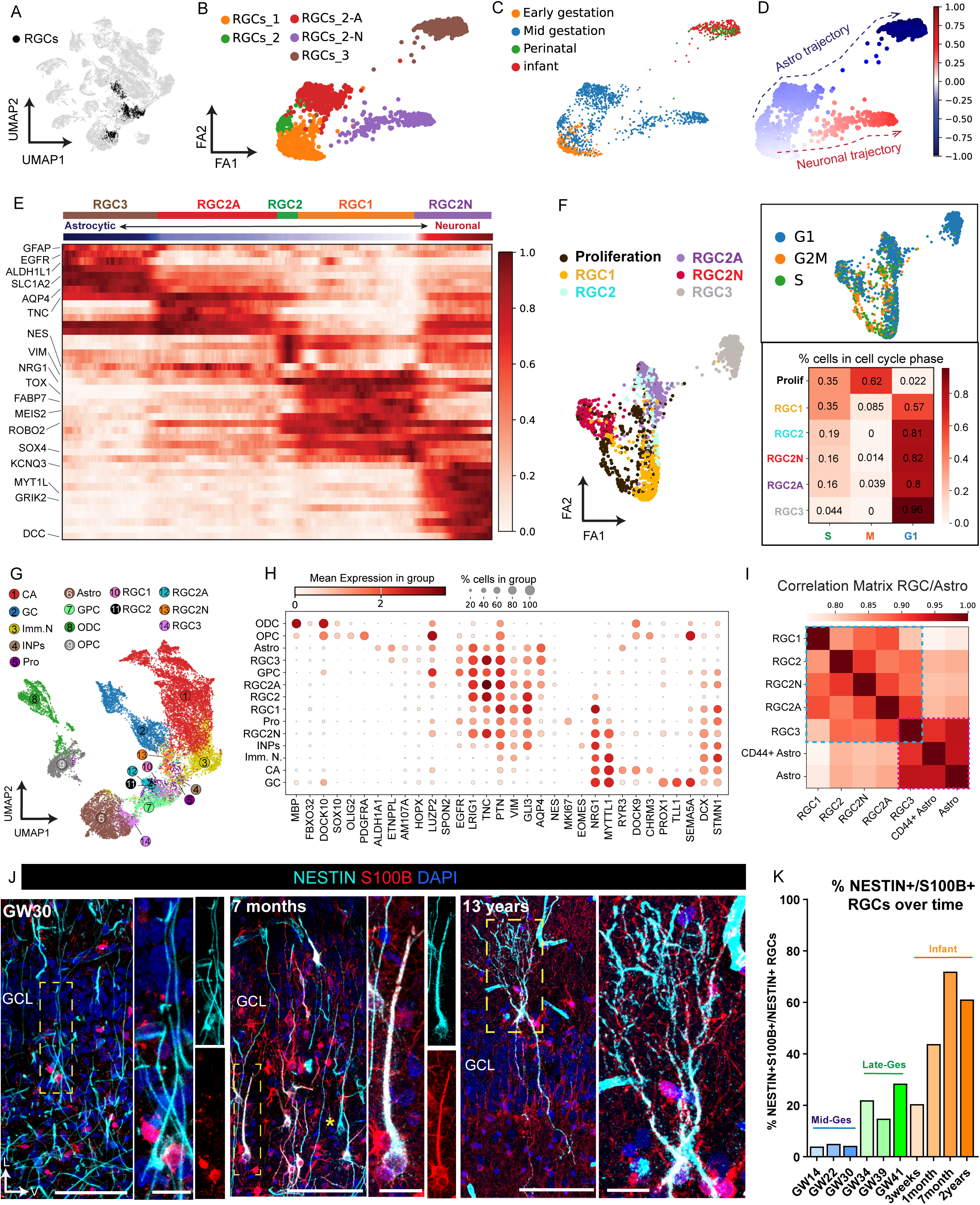
Transcriptomic and histological characterization of radial glial cell developmental progression in the human dentate gyrus. **(A)** UMAP embedding with cells classified as RGCs highlighted in black. **(B-D)** UMAP embedding of RGCs annotated by RGC subclasses, identified through unsupervised clustering and manual annotation (B), developmental stage (C), and pseudotime trajectory from an RGC1 root cell (D). **(E)** Heatmap showing top markers defining the pseudotime trajectory described in C. **(F)** Left: UMAP embedding of identified RGC subtypes and proliferating cells. Right: UMAP embedding of cell cycle score analysis identifying cells in G1, G2M or S phase (top) and percentage of cells on each cell cycle phase per cell subclass (bottom). **(G)** UMAP embedding of RGC subtypes together with major excitatory neurons, oligodendrocyte and astrocyte lineage cells. **(H)** Dotplot for markers defining the identity of each subtype from G. **(I)** Correlation matrix between RGC subtypes and the two major astrocyte subtypes. **(J)** Immunofluorescence for NESTIN and S100B in the GCL and GHPZ region at GW30, 7 months and 13 years. **(K)** Quantifications showing the percentage of radial cells co-expressing NESTIN and S100B in the GCL and GHPZ in ten individual cases from GW14 to 2 years. The data is shown as the mean of 2 to 5 sections from distinct anterior–posterior levels per case. Each age is represented by a n=1 case. <u>Abbreviations</u>: CA, cornu ammonis; FA, force atlas; GC, granule cell; GCL, granule cell layer; GHPZ, granular-hilar progenitor zone; GPC, glial progenitor cell; GW, gestational week; INP, intermediate neural progenitor; L, lateral; ODC, oligodendrocyte; OPC, oligodendrocyte progenitor cell; RGC, radial glia cell; UMAP, uniform manifold approximation and projection; V, ventral. <u>Scale bars:</u> (J) 100 µm and 20 µm (High magnification).

Motivated by the lack of RGC signatures in our adult sample, we further investigated the presence of RGC signatures in the postnatal period. We first integrated our dataset with infant and early childhood cases (0-5 years) from a published human hippocampal dataset (∼135,000 cells)^24^ and manually annotated the cells based on unbiased clustering with our cell populations. Together with major neuronal and glial populations, a substantial number of cells from the Dumitru et al. dataset (3.30% of the dataset; 4,401 cells) grouped with a combined cluster of RGC2As and RGC3s from our dataset, confirming the strong presence of RGCs in the infant period. Interestingly, 1.26% of cells (1,681) formed a distinct cluster with RGC2Ns and appeared located between inhibitory neurons and OPCs in the UMAP space. Next, we projected cells from the adult cases of the same hippocampal dataset (∼363,000 cells, 13-78 years) into the integrated developmental UMAP, observing changes in cell type composition. Only 0.04% of cells (134 cells) were assigned an RGC2A-RGC3 identity, and the RGC2N group was also reduced, with 0.21% of cells (770 cells) assigned to this identity. Of note, several cells were classified as immature neurons of both CA and granule neuron identity, supporting the presence of neurons at different maturation states in both the infant and adult period **(Figure S5A)**. We further investigated the identity of cells classified as RGCs and excitatory neurons using canonical markers. Cells categorized as RGC2A-RGC3 in the infant period showed lower GFAP expression than cells categorized as astrocytes, but higher expression of NESTIN and the WNT-associated genes WNT7B and LEF1. Moreover, DCX and PROX1 were also expressed, confirming the granule neuron-bound fate of these cells. In contrast to these observations, RGC2A–RGC3s from adult cases showed high GFAP expression and a complete lack of NESTIN, LEF1, WNT7B, DCX, or PROX1, revealing that the few cells identified as RGC2A-RGC3 displayed a strong astrocytic identity. Cells classified as RGC2Ns showed strong GFAP, DCX, PROX1 and also GAD1 expression in both infant and adult cases, further supporting an inhibitory neuronal nature of this population. We observed DCX expression in every excitatory neuron cell type from infant cases, and in cells that mapped onto CA3 and CA3 immature neurons from the adult cases. Interestingly, cells classified as CA3 immature neurons from both infant and adult period showed GAD1 expression, suggesting inhibitory identity **(Figure S5B)**. These results highlight the lack of RGC identities in the adult DG, as observed during infancy and early childhood.

We next defined the molecular progression of the observed RGC subtypes. Pseudotime ordering revealed a bifurcating trajectory originating from RGC1 **(Figure 5D)**. Heatmap illustrating the top five genes per RGC subtype defining pseudotime progression revealed VIMENTIN, FABP7 and TOX as RGC markers specifically enriched in RGC1 compared to the rest of RGCs. Moreover, genes associated with immature neurons, such as NRG1, ROBO2 and SOX4, further defined this RGC subtype. MEIS2, previously shown to be associated with CA, also showed enrichment in RGC1 and the smaller transition subtype RGC2, suggesting presence of CA-bound progenitors within these subtypes. From RGC1, we observed one trajectory gradually acquiring genes associated with an astrocytic identity, including SLC1A2 and ALDH1L1, alongside increasing AQP4 and GFAP expression (the RGC2A → RGC3 trajectory). The canonical RGC markers NESTIN and TNC remained highly expressed in these groups, supporting their RGC identity. The second trajectory, defined as RGC2N, progressed toward a neuronal fate, acquiring expression of neuron-related genes such as KCNQ3, MYT1L, GRIK2 and DCC **(Figure 5E)**. Molecular distinctions among RGC subtypes were also observed by cell cycle scoring analysis. Proliferative activity was highest in RGC1 (35% of cells in S phase, 8.5% in M-phase), progressively decreased in RGC2, RGC2A and RGC2N (16-19% of cells in S-phase, 0-3.9% in M-phase), and was markedly reduced in RGC3 (4.4% in S-phase and 0% in M-phase) **(Figure 5F)**. These analyses support the importance of mid-gestation as a time window during which lineage-committed RGC subtypes emerge (RGC2A and RGC2N), highlighting the postnatal persistence of a highly quiescent subtype that acquires astrocytic identity (RGC3).

We sought to further define the transcriptomic identity of each RGC subtype by reclustering them with OPCs, oligodendrocytes, astrocytes, granule neurons and CA neurons. In this integrated embedding, RGC2N clustered adjacent to the granule cell lineage, whereas RGC3 was positioned within the astrocyte cluster **(Figure 5G)**. Canonical marker expression supported these relationships: The RGC markers TNC, VIMENTIN, GLI3 and NESTIN, although at varying degrees, showed enrichment in all RGC subtypes. However, the infant subtype RGC3 showed expression of the astrocyte markers ALDH1L1, ETNPPL and S100B, while the rest of the RGCs did not. Moreover, DCX and STMN1, canonical immature neuron markers, were expressed in RGC1, RGC2N and RGC2A but were absent in RGC3s **(Figure 5H)**. Correlation analysis of transcriptomic similarity between RGC subtypes and the two major astrocyte subtypes corroborated the astrocytic nature of RGC3, which displayed higher similarity scores to astrocytes than the rest of RGC clusters. (**Figure 5I**). Collectively, these analyses confirm RGC3, the infant RGC population, as a subtype with emerging astrocytic features while retaining a core RGC identity. Collectively, these analyses highlight RGC3, the infant RGC population, as a subpopulation at the interface of RGC and astrocytic identity, potentially representing a transitional state with emerging astrocytic features or an early astrocyte subtype retaining core RGC markers.

To understand the observed transcriptomic trajectories within a cellular context, we used NESTIN and S100B expression to label RGCs and astrocytes, respectively, in the human DG from mid-gestation to adulthood. At GW22 through GW34, NESTIN^+^ radial cells occupying the GHPZ and GCL were clearly distinguishable from S100B^+^ astrocytes in the same territory, with rare co-expression of both markers. After birth, NESTIN^+^ radial cells were still present in the GHPZ and exhibited similar radial morphology to that seen at earlier stages, with their soma located in the GHPZ and a single radial process extending toward the molecular layer of the DG. However, S100B^+^ astrocytes with stellate morphology increased across the region and a substantial portion of NESTIN^+^ cells also co-expressed S100B, despite conserving a radial morphology. After 7 years, the rare NESTIN^+^ cells observed across different regions of the DG exhibited a multipolar morphology with multipolar branching and co-expressed S100B (**Figures 5J and S5C**). We quantified NESTIN^+^ radial cells, defined as cells with a single radial process extended toward the molecular layer and their nucleus located within the GHPZ and granule layer, and their co-expression with S100B from early gestation to infancy. The results showed less than 10% of NESTIN^+^ radial cells co-expressed S100B before GW30, but the presence of double positive cells gradually increased over time. After 7 months, more than half of radial NESTIN^+^ cells were S100B^+^, consistent with a gradual differentiation toward an astrocytic phenotype (**Figure 5K; Table S2)**. Additionally, we tested the RGC marker VIMENTIN to interrogate its expression patterns. At GW37, although VIMENTIN^+^/NESTIN^+^ radial cells were present in the GCL and GHPZ, VIMENTIN expression was also observed in S100B^+^ cells with stellate morphology in different regions **(Figure S5D**), highlighting widespread expression of VIMENTIN in RGCs and astrocytes during the gestational period. Altogether, these results support the transcriptomic trajectory observed before and confirm the transition of RGCs toward an astrocytic phenotype during infancy.

### WNT signaling defines NPC identity

Motivated by the observed developmental transition of NPCs, we interrogated the molecular regulators driving transcriptomic remodeling. Differential gene expression analysis yielded broad RGC -specific upregulated gene sets. RGC2 showed the fewest uniquely enriched genes (298), followed by RGC2A (409) and RGC1 (1298). Consistent with a more differentiated identity, RGC2N and RGC3 showed a bigger set of uniquely upregulated genes (adjusted p < 0.01, log2 fold change > 0.5), with 2745 and 1425, respectively **(Figure 6A)**. Evaluation of the top upregulated genes in each RGC subset revealed genes associated with WNT signaling. RSPO3, LEF1 and WNT7B were significantly increased in RGC2A, while WIF1 was enriched in RGC3, suggesting a role for WNT in this trajectory of RGCs **(Figure 6B)**. We performed pathway enrichment analysis (KEGG and Reactome) on each of these gene sets, identifying significantly upregulated signaling pathways (adjusted p < 0.05). The results confirmed the activation of WNT signaling in RGC2A and, to a lesser extent, in RGC3, supporting WNT signaling as a key pathway driving RGC maturation toward postnatal stages. Notably, ErbB pathway was upregulated in the RGC2N subtype and Hippo and Notch pathways, previously associated with cell quiescence and regulation of adult hippocampal neurogenesis^48–50^, were activated in RGC3s **(Figure 6C)**. Individual evaluation of the genes contributing to pathway activation confirmed that LEF1 and WNT7B, previously identified as markers of RGCs in the GHPZ (Figure 3), were strongly upregulated in RGC2A, consistent with the WNT signaling signature of this subtype. In contrast, the WNT-related genes like TLE1, TLE3, SMAD3, BAMBI or WIF1 and NOTCH2 showed expression in both RGC3s and astrocytes, consistent with shared genetic programs between these subtypes **(Figure S6A; Table S7)**. These data highlight specific pathways associated with each RGC subtype observed across the developmental trajectory, suggesting WNT signaling pathway as a major regulator of RGC identity maintenance.

**Figure 6.**
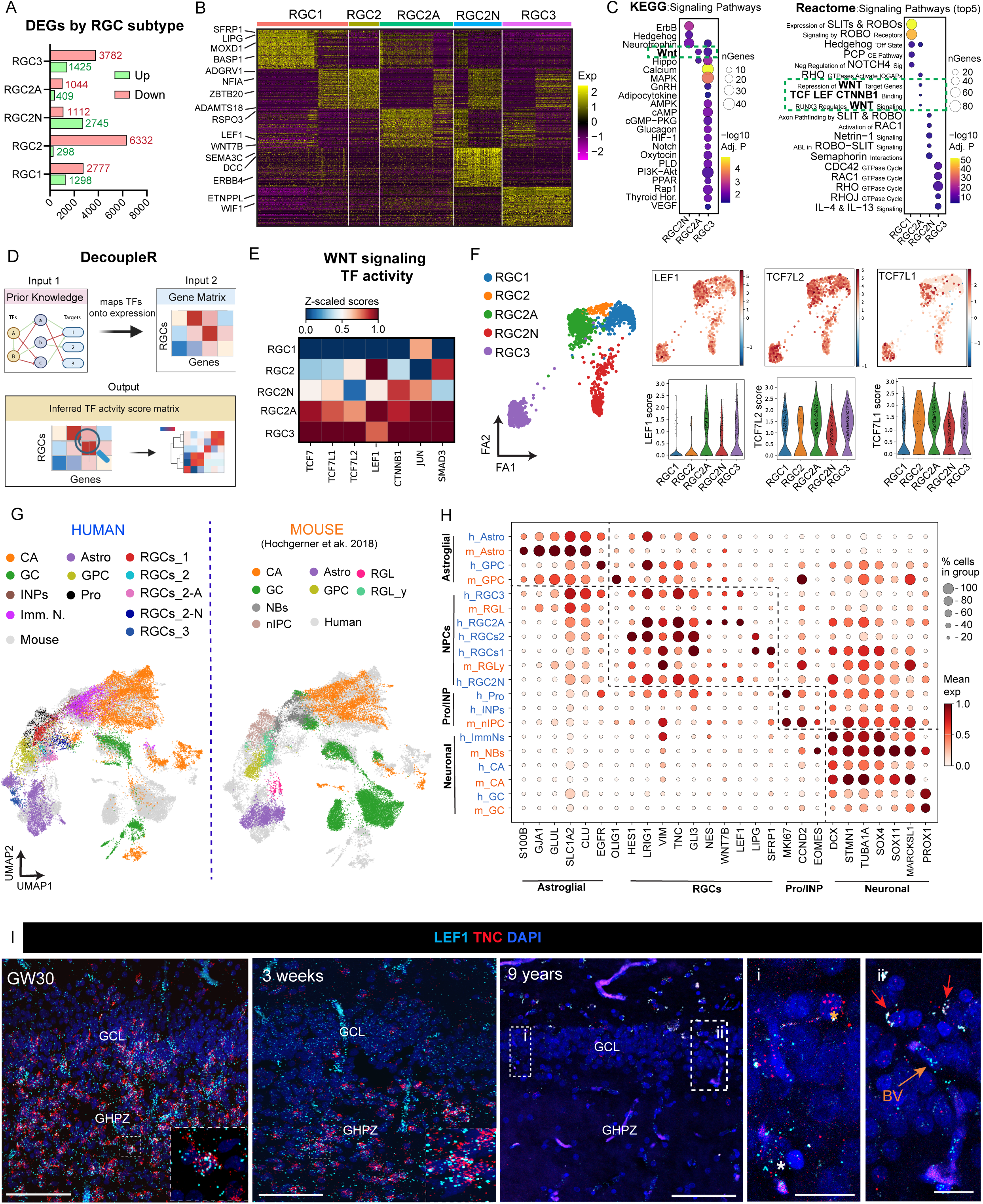
Identification of signaling pathways defining RGC subtypes in the developing DG. **(A)** Bar graph showing differentially upregulated (green) and downregulated (red) genes in each RGC subtype (adjusted p < 0.01, Log₂FC > 0.5). **(B)** Heatmap showing the top 100 upregulated genes per RGC subtype. Neurodevelopmentally relevant genes are highlighted. **(C)** Dot plot of signaling pathway enrichment by RGC subtype (KEGG 2024). Right: Dotplot of top 5 enriched signaling pathways per RGC subtype (Reactome 2024). **(D)** Schematic of the decoupleR workflow applied to RGC subtype DEGs to infer TF activity. **(E)** Heatmap of WNT-associated TFs with enriched inferred activity across RGC subtypes identified by decoupleR. **(F)** UMAP embedding generated from the decoupleR pipeline showing RGC subtype distribution, with inferred TF activity scores for LEF1, TCF7L2, and TCF7L1 projected onto the UMAP. Violin plots show the distribution of inferred activity scores per RGC subtype. **(G)** UMAP embedding showing integration of the human snRNA-seq dataset with the mouse hippocampal dataset (Hochgerner et al., 2018). Selected cell types, including excitatory neurons (CA and GC), intermediate progenitor cells (INPs), immature neurons, astrocytes, glial progenitor cells (GPCs), proliferating cells, and RGC subtypes are annotated across both datasets. Human (left) and mouse (right) datasets are shown separately. **(H)** Dotplot with canonical markers for the cell types shown in G. Cell types are shown separately for humans (blue) and mice (orange). **(I)** smFISH for LEF1 and TNC in the GCL and GHPZ regions at GW30, 3weeks and 9 years. Square boxes in the 9 year picture show high magnification images of a cell co-expressing LEF1 and TNC (white asterisk) and a TNC-positive cell near the molecular layer (yellow asterisk) (i) and a blood vessel expressing LEF1 (orange arrow)(ii). Red arrows signal lipofuscin aggregates. <u>Abbreviations:</u> CA, cornu ammonis; D, dorsal; FA, Force atlas; GC, granule cell; GCL, granule cell layer; GHPZ, granular-hilar progenitor zone; GPC, glia progenitor cell; GW, gestational week; INP, intermediate neural progenitor; L, lateral; NB, neuroblast; nIPC, neural intermediate progenitor cell; RGC, radial glia cell; RGL, radial glia like; RGL_y, Radial glia like young; UMAP, uniform manifold approximation and projection. <u>Scale bars:</u> (I) 100 µm and 20 µm (High magnification).

To functionally contextualize these transcriptional differences, we applied DecoupleR using curated regulons, enabling quantitative estimation of transcription factor activation scores across RGC subtypes^51^ **(Figure 6D)**. Consistent with the observed role of the WNT pathway in RGCs, LEF1 was significantly active in RGC2A **(Figure S6B)**. Moreover, the core effectors of canonical WNT/β-catenin signaling TCF7, TCF7L1, TCF7L2 and CTNNB1, and other transcription factors associated with WNT signaling activity, such as JUN and SMAD3, showed significant activation scores in RGC2A and RGC3. Notably, LEF1 displayed higher activity in RGC2A relative to RGC3 **(Figure 6E)**, suggesting a decrease in WNT signaling modulation through this effector over time. The differential activity patterns were further corroborated by the heterogeneous score distributions observed in the UMAP embedding and violin plot representations **(Figures 6F)**. These findings support the notion that RGC2A and RGC3 are distinguished not only by WNT-associated marker gene expression but also by divergent upstream transcriptional regulatory states, as corroborated by elevated inferred activity scores for WNT-related transcription factors in these subtypes.

To evaluate the conservation of the identified human RGC subtypes across species, we integrated our dataset with a previously published mouse single-cell RNA sequencing dataset^52^, which included RGCs from the murine embryonic day 16 to postnatal day 5 (radial glia young, RGLy) and adult stage (radial glia-like, RGL). UMAP visualization of major neurogenesis-related populations showed expected clustering of equivalent cell populations, with two distinct pseudotime trajectories from the youngest human and mouse RGCs toward neuronal and astrocytic cell types. Interestingly, human prenatal RGC subtypes were closely grouped with the murine developmental RGLy subtype, whereas human RGC3s and mouse RGLs were more proximal to astrocytes **(Figure 6G and S6C)**. The trajectory of RGCs toward astrocytes in both mouse and human was supported by pseudotime analysis of the combined RGC subtypes, GPCs and astrocytes **(Figure S6D)** and dot plot visualization of canonical marker genes confirmed transcriptional correspondence between murine RGL and human RGC3. Both subtypes exhibited increased expression of astrocyte markers (GJA1, GLUL, SLC1A2 and CLU) and decreased expression of immature neuron markers (DCX, STMN1, TUBA1A, SOX4, SOX11 and MARCKSL1) compared to RGCs from younger stages. Nonetheless, ALDH1L1 and S100B, highly expressed in human and mouse astrocytes, were also expressed in human RGC3s, suggesting an even stronger astrocytic identity for this subtype. Despite the more differentiated signature of RGC3 and RGL, all RGC groups in both human and mouse expressed canonical RGC markers, such as HES1, VIMENTIN, TNC and NESTIN. The previously identified RGC1 marker SFRP1 was also observed in murine RGs, whereas LIPG showed restricted expression to human RGC1. Importantly, LEF1 and WNT7B, expressed in human RGC2A and RGC3 with a decreasing trend over development, showed consistent expression in murine RGCs across both developmental and adult stages **(Figure 6H)**. These data reveal a temporal progression toward an astrocytic transcriptomic phenotype in both mouse and human RGCs, and highlight a species difference in temporal regulation of LEF1 and WNT7B.

To validate LEF1 and WNT7B expression in the anatomical context, we spatially mapped these markers in the developing human DG using single-molecule fluorescence in situ hybridization (smFISH). At GW30 we observed abundant LEF1^+^ cells in the GHPZ, colocalizing with the RGC marker TNC. Moreover, from GW18 to GW37, the majority of LEF1^+^ cells within the DG co-expressed WNT7B, but not the vascular marker CD31, confirming LEF1 and WNT7B as DG-associated RGC markers. LEF1^+^ cells that co-expressed TNC and WNT7B were still observed in the GHPZ at 3 weeks after birth. However, the expression of LEF1 and WNT7B in the GHPZ declined during childhood. At 9 years, WNT7B⁺ cells were rare and LEF1 expression was largely vascular, paralleling the above-described disappearance of NPCs at this stage **(Figures 6I and S6E)**. No cells positive for both LEF1 and WNT7B were observed around the GCL at this age. Large-sized puncta appearing double-positive for smFISH probes were identified as lipofuscin deposit autofluorescence, present in the 9 year-old human tissue sections across all fluorescent channels **(Figures 6I and S6E)**. Together, these data confirm LEF1 and WNT7B as defining molecular features of NPCs in the human DG across early prenatal to postnatal stages, with developmental expression trajectories that parallel the loss of NPCs in the human DG by childhood.

## Discussion

Here, we generated a multi-modal atlas that integrates morphological characteristics, spatial coordinates, and transcriptomic identity of NPCs across the pre- and postnatal human DG. We demonstrate that NPCs converge within the DG by mid-gestation, forming a neurogenic niche that we termed the GHPZ. We demonstrate that RGCs persist in the GHPZ in the infant period, progressively transitioning toward an astrocytic fate, with only a sparse population remaining into childhood. We further propose LEF1 and WNT7B as molecular markers for the identification of GHPZ-associated RGCs during the prenatal and early postnatal period. Our findings deepen our understanding of NPC development in the human DG and establish a reference framework to reconcile discrepancies in NPC identification in the adult human brain.

### The unique timeline of NPC organization in the developing human DG

Our work builds on prior studies^16,18^ and delivers an integrated cellular and molecular characterization of the transient neurogenic niches that drive the formation of the human DG (the DVZ and the DMS). The cellular composition of the DVZ-DMS, containing NESTIN^+^ RGCs, TBR2^+^ INPs and PROX1^+^ cells that define granule neuron identity, is comparable to the cytoarchitecture described in rodents^15^. At the temporal scale, we observed maturation of the DVZ and progressive depletion of the DMS during the mid-gestational period. This coincides with the rapid angular reorientation of the hippocampus from a dorsal-ventral to a rostral-caudal configuration, a key morphogenetic transition in human hippocampal development. These observations differ markedly from the persistence of the DMS until at least postnatal day five in rodents^10,11,53^ and demonstrate the relative compression of DG formation into an earlier relative developmental window in humans, as previously proposed^54,55^. Furthermore, this study confirms previous observations that highlighted the lack of a rodent SGZ-like NPC organization in the human DG^18^. Instead, a diffuse multilayer territory containing RGCs, INPs and migratory neurons and enriched for WNT-associated genes is established between the hilus and the GCL by mid-gestation, herein termed the granular hilar progenitor zone (GHPZ). Importantly, while the rodent SGZ is maintained into adulthood, the GHPZ in humans disappears by childhood, consistent with an overall accelerated NPC depletion timeline in humans. One potential explanation for these temporal discrepancies is the absence of a species-specific mechanism to establish a lifelong NPC pool. In rodents, adult RGCs emerge postnatally through a cyclin D2-dependent process^56^. Notably, our transcriptomic analysis reveals a marked reduction in cyclin D2 expression in human RGCs compared to their mouse counterparts, suggesting that this expansion mechanism may not be conserved across species.

### Definition of RGCs in the human DG

The absence of standardized molecular and morphological criteria for identifying RGCs has hampered efforts to develop a cohesive perspective on the extent of RGC presence in the human DG. Building on criteria established in rodent studies, we defined DG-associated RGCs as cells with their soma located in the GHPZ, a single radial process extended toward the molecular layer, and expression of the molecular marker NESTIN^31^. Moreover, our transcriptomic analysis identified LEF1 and WNT7B as additional markers for RGCs in the human DG. LEF1 has been extensively reported as a marker of RGCs in the rodent DG, whereas WNT7B remains largely unexplored. Importantly, the identification of RGCs is further complicated by heterogeneity within the RGC population itself. We expanded on prior studies that described distinct progenitor subtypes in the human hippocampus between GW16 and GW27^23^, and identified two transcriptomically distinct RGC subtypes at mid-gestation, with neuronal and glial biases, respectively, and differences in cellular organization and temporal progression between the medial and lateral blades of the DG. This heterogeneity converged in an overall trajectory of RGCs toward an astrocytic phenotype and decline in numbers, shared between mouse and human. However, while RGC heterogeneity encompassing different quiescent and activated states and self-renewal capacities has been documented in the adult rodent DG ^57,58^, whether analogous RGC subtypes with distinct cellular behaviors co-exist in the postnatal human DG remains an open question. Our cross-species analysis suggests transcriptomic equivalence between adult mouse and early postnatal human RGCs, yet deeper transcriptomic and morphological characterization will be required to fully resolve this question.

### From NPCs to astrocytes

In the adult rodent DG, NPCs differentiate into astrocytes following neurogenic divisions, leading to a decrease in neurogenic capacity over time^57,59^. Consistent with this progressive shift, remaining NESTIN^+^ radial cells in the aged SGZ acquire astrocytic properties, such as S100B expression, and exhibit lower proliferative potential^60^, reflecting a niche that becomes increasingly astrogenic with age^61^. Here, we reveal an equivalent progression of NPCs in the developing human DG, albeit compressed into an earlier developmental time window between the third gestational trimester and infancy. Whether the transcriptomic shift of RGCs toward an astrocytic phenotype reflects a true gliogenic switch, analogous to what has been described in the human neocortex^47,62^, or whether these cells retain neurogenic capacity despite this transcriptomic transition, remains to be determined. Indeed, neurogenic and gliogenic programs have been shown to overlap in the rodent hippocampus^63^, and our data suggest a similar coexistence during human DG development. Expression of LEF1, reported to promote the expression of the granule neuron marker PROX1^23^, suggests that infant NPCs retain neurogenic capacity. In contrast, several converging lines of evidence indicate a reduction in neurogenic potential and an increase in gliogenic activity across this progression. Changes in WNT signaling, including downregulation of its effector LEF1 and upregulation of its inhibitor WIF1, previously associated with progenitor quiescence^64^, suggest a weakening of the pro-neurogenic WNT cascade by infancy. Concurrently, enrichment of NFIA and SOX9^65^ points toward an active gliogenic program, consistent with their established roles in suppressing neurogenesis while promoting astrocytic fate. Further supporting a shift toward quiescence, infant NPCs show enrichment of genes associated with HIPPO, AMPK, NOTCH, and STAT signaling — pathways collectively linked to neurogenic decline^49,66^. Together, these transcriptomic signatures suggest that the NPC progression toward a glial phenotype is driven by the coordinated activation of multiple gene regulatory networks. Future investigation of NPC expansion and differentiation dynamics using *in vitro* platforms could help define the molecular programs underlying NPC lineage progression.

### Brain plasticity beyond NPCs

The cellular complexity of the human brain is largely built on prolonged neurogenesis^67–70^. However, our data reveal that the DG is a notable exception, with a compressed window compared to rodents. Neurogenesis, understood as the process by which an NPC divides and generates a newborn neuron that ultimately integrates into the neuronal circuitry, has been proposed to contribute to forgetting and to disrupt the microenvironment of existing neurons^71,72^. Therefore, one possibility is that NPC-driven neurogenesis in the human DG has been replaced by a more stable system of protracted neuronal maturation through evolution, consistent with neoteny observed in other human brain areas^73,74^. Recent transcriptomic studies have reported extensive numbers of granule neurons showing expression of genes associated with immature properties^20,75,76^. Our results support the presence of immature neurons in the adult hippocampus and provide a developmental perspective, showing immature neurons in the CA layer during the third gestational trimester and inhibitory migratory currents that appear by mid-gestation. Furthermore, immature genes, such as DCX, were present in CA and inhibitory lineages beyond gestation into postnatal life. While the precise maturation time course of these populations remains to be determined, we postulate that neuronal neoteny is a broader feature of hippocampal circuitry beyond the DG.

Finally, our findings underscore the presence of NPCs persisting into the infant period in the human DG, potentially analogous to the adult NPC reservoir characterized in rodents. The responsiveness of this progenitor pool to pathological insults is supported by reports of increased Nestin-expressing cells in the infant human hippocampus following seizures^77^, and an extensive rodent literature documenting NPC activation in traumatic brain injury, epilepsy, and ischemia^78–80^. The existence of a comparable progenitor population in the infant human hippocampus, which can be exposed to perinatal and childhood injuries, thus opens a compelling translational avenue, inviting direct investigation of NPC-mediated responses in human hippocampal pathology.

## Acknowledgements

This work was supported by the National Institutes of Health (grant P01 NS083513 to M.F.P. and U01MH130995, to C.L. and M.F.P.); (grant R01MH125252 and RF1MH130461 to C.L..); the Roberta and Oscar Gregory Endowment in Stroke and Brain Research (to M.F.P.); the Chan Zuckerberg Initiative (to M.F.P.); and the National Science Foundation under CAREER Award 2337373 (to J.G). Confocal microscopy images were acquired at the UCSF Innovation Core at the Weill Institute for Neurosciences. The bulk-RNA spatial transcriptomics experiment using GeoMx technology was performed at the Laboratory for Cell Analysis at the UCSF Hellen Diller Family Comprehensive Cancer Center. We thank Kenneth X. Probst for illustration.

## Data and code availability

Fastq files and processed single-nucleus RNA data are available through the NeMO archive under accession nemo:dat-ortissu https://assets.nemoarchive.org/dat-ortissu

An interactive browser to visualize CosMx single cell spatial transcriptomics data is available at https://hpc.igvfspatial.com/

## Materials and Methods

### Brain Specimen Collection

Post-mortem human brain tissue was collected at the University of California, San Francisco (UCSF) with post-mortem intervals (PMI) of less than 24 hours, under prior patient consent and in accordance with the ethical regulations of the UCSF Committee on Human Research. All protocols were approved by the Human Gamete, Embryo and Stem Cell Research Committee (IRB GESCR# 10-02693). Tissue from gynecology clinics was collected from patients who expressed interest in donating tissue for research purposes and provided written informed consent following both written and oral information. Gestational age was estimated using clinical data including last menstrual period, ultrasound measurements, crown-rump length, and anatomical landmarks. All specimens were evaluated by a neuropathologist as control samples through the UCSF Pediatric Neuropathology Research Laboratory. Additional samples used for electron microscopy were provided by Biobank La Fe (B.0000723) and handled in accordance with Spanish Royal Decree 1716/2011 governing the use of human biological samples in biomedical research. This component of the study was approved by the corresponding Institutional Ethics Committee (2018/0356) and Scientific Committee, and written informed consent was obtained from all donors through the Neonatology and Gynecology Units of Hospital Universitari i Politècnic La Fe. Postnatal samples from individuals with a history of neurological disorders or trauma were excluded from primary analyses. Prenatal samples with abnormal neuropathology or positive for common chromosomal aberrations were similarly excluded. Two samples that did not meet these primary inclusion criteria, one with a chromosome 1 deletion and one with a history of hypoxic-ischemic injury, were retained for supplemental analyses **(Figure S1I and S2F)**, where findings are presented descriptively and interpreted with appropriate caution. The metadata of the cases used in this study can be found in **Table S1**.

Tissue was sectioned coronally and regions of interest were dissected. For single-cell RNA sequencing, 1 mm tissue blocks were flash-frozen in liquid nitrogen and stored at −80°C until use. Blocks designated for histological and spatial transcriptomic analyses were fixed in 4% paraformaldehyde (PFA) for 48 hours, cryoprotected in 30% sucrose, embedded in OCT, and cryosectioned onto glass slides. For each case, three cresyl violet-stained sections spanning the block were examined to confirm anatomical positioning using established landmarks. For transmission electron microscopy (TEM), one hemisphere was processed by immersion fixation for 7 days; tissue designated exclusively for ultrastructural analysis was fixed in 2% PFA and 2.5% glutaraldehyde, while samples intended for pre-embedding immunogold procedures were fixed in 4% PFA.

### Structural MRI of Fetal Hippocampus Development

To gain an understanding of the development of the hippocampus macrostructural features, such as its orientation and gross volume, we analyzed a neuroimaging dataset of T2-weighted fetal MRI volumes spanning the gestational period of 21 to 38 weeks. As described in its initial release^28^, a T2-weighted SSFSE protocol was employed with 0.9mm isotropic in-plane resolution, 2mm slice thickness, TE = 110-120ms, and TR = 1400-2000ms, with a 3-Tesla Siemens Skyra. At least 6, but up to 23 fetuses, were used to construct an average brain template at each week of gestation (21 to 38 GW). Imag s were aligned to the plane formed by the anterior and posterior commissures (AC-PC) and resampled to 800 micrometer isotropic voxels. Brain volumes were segmented based on anatomical borders[cite], from which the hippocampal label was extracted for further analysis. In addition to the overall volume of the hippocampus at each time point, we quantified the orientation of the hippocampus by measuring the angle formed by the long axis of the hippocampus with the vector denoting the AC and PC points. To this end, two coordinates were identified in each hippocampus: one in the posterior-most aspect of the hippocampal tail, and one in the anterior-most ported of the hippocampal head (see Figure 1 for examples). The dot product (arc cosine) between this vector and that of the AC-PC vector was derived and converted to degrees. To quantify how the hippocampal angle potentially changes with development, we modeled the hippocampal angle as a function of gestational age using an exponential decay model defined as:

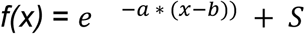

Where ***a*** describes the steepness of the decay, ***b*** captures horizontal shift of the function, and ***S*** captures the vertical shift of the function. The model was fit by optimizing parameters through a minimization of root-mean-square-error, with final fit values of a = 0.116, b = 50.75, S = 33.63, and a model-fit R^2^ = 0.93. To further identify the inflection point when the slope of this decay significantly changes from initial steep decline to a smaller decay, we performed a two-lines test which, after removing a linear fit from the data, tries to identify the x-value (gestational age) at which two lines of different slope can be fit. This test finds that the best-fitting point of two lines occurs at 25 gestational weeks, with the data prior to this timepoint having a slope significantly different from later timepoints (slopes = −4.83 and 0.72, p’s < 0.001).

### Immunofluorescence Staining

Tissue sections were cryosectioned at 30 µm, mounted onto Superfrost Plus slides, and stored at −80°C until use. Prior to staining, slides were baked at 65°C overnight, then rehydrated in TNT buffer (PBS containing 0.05% Triton X-100), and post-fixed in 4% paraformaldehyde for 20 minutes. Endogenous peroxidase activity was quenched by incubation in 1% H₂O₂ in PBS for 30 minutes. Where applicable, antigen retrieval was performed by heating slides in a 10 mM sodium citrate buffer (pH 6.0) at 95–100°C for 10 minutes, followed by cooling at room temperature for 30 minutes. Sections were then incubated again in 1% H₂O₂ in PBS for 1 hour, and subsequently blocked for 1 hour in TNB solution (0.1 M Tris–HCl, pH 7.5; 0.15 M NaCl; 0.5% blocking reagent, PerkinElmer). To improve signal amplification, postnatal cases older than 2 years were stained following the same protocol but with a stronger TNT buffer (PBS containing 0.2% Triton X-100) and TNB solution (0.1 M Tris–HCl, pH 7.5; 0.15 M NaCl; 2.0% blocking reagent, PerkinElmer). Both protocols were validated in tissue from donors younger than 2 years, with no differences observed between conditions.

Primary antibodies diluted in TNB were applied overnight at 4°C. The following day, sections were incubated with either biotinylated secondary antibodies (1:250 in TNB) or Alexa Fluor–conjugated secondary antibodies, together with DAPI to label nuclei, for 2.5 hours at room temperature. For tyramide signal amplification (TSA), biotinylated secondary antibody incubation was followed by streptavidin-HRP (1:200 in TNB) for 30 minutes, and subsequently by tyramide-conjugated fluorophores diluted in amplification buffer (PerkinElmer) in the following order: Cy5 (1:100), fluorescein (1:50), and Cy3 (1:100). Sections were washed with a TNT buffer between all steps. Following staining, sections were dehydrated, mounted with slide mounting medium, and coverslipped.

| Antibody | Host/Isotype | Dilution | Supplier | Catalog number |
| --- | --- | --- | --- | --- |
| Anti-Nestin | Mouse | 1:250 | Abcam | ab18102 |
| Anti-S100B | Rabbit | 1:500 | Novus | NBP2-54426AF594 |
| Anti-TBR2 | Sheep | 1:200 | ThermoFisher | PA547818 |
| Anti-Ki67 | Mouse | 1:250 | BD Pharmingen | 556003 |
| Anti-DCX | Rabbit | 1:250 | Cell Signaling | 4604S |
| Anti-PROX1 | Goat | 1:200 | R&D | AF2727 |
| Anti-SOX2 | Goat | 1:250 | SCBT | sc-17320 |
| Anti-HOPX | Rabbit | 1:250 | Proteintech | 11419-1-AP |
| Anti-GFAP | Chicken | 1:500 | Abcam | ab4674 |
| Anti-Vimentin | Chicken | 1:250 | Invitrogen | PA1-10003 |
| Anti-GAD67 | Mouse | 1:200 | SCBT | sc-28376 |
| Anti-CD31 | Sheep | 1:250 | R&D Systems | AF806 |

### Transmission Electron Microscopy

Samples were initially sectioned into 5 mm-thick coronal slices in order to determine at what level of the anteroposterior axis we are located. Once the region of interest was selected, it was cut at 200 or 100 µm using a vibratome VT-1000S (Leica Microsystems, Wetzlar, Germany). Sections were post-fixed in 2% osmium tetroxide (Electron Microscopy Sciences, Hatfield, PA, USA) for 1.5 hours, rinsed, dehydration initiated, contrasted with 2% uranyl acetate (Electron Microscopy Sciences, Hatfield, PA, USA) in 70% ethanol for 2.5 hours, completely dehydrated through a graded series of ethanol, and embedded in Durcupan resin (Fluka, Buchs, Switzerland).

Semithin sections (1.5 µm thick) were obtained using an UC-6 ultramicrotome (Leica, Heidelberg, Germany), mounted on gelatin-coated glass slides, and stained with 1% toluidine blue (Panreac, Barcelona, Spain). Sections were examined under a light microscope Eclipse E800 (Nikon, Tokyo, Japan).

For identification of different cell types and their ultrastructural characteristics, ultrathin sections (60–70 nm) were cut using the same ultramicrotome, stained with lead citrate (Reynolds solution), and examined with a transmission electron microscope Tecnai Spirit G2 (FEI, Eindhoven, The Netherlands). Digital images were acquired using a XAROSA digital camera (EMSIS GmbH, Münster, Germany) controlled by Radius software (version 2.1).

### Fluorescence In Situ Hybridization (smFISH)

Tissue sections were cryosectioned at 30 µm, mounted onto Superfrost Plus slides, and stored at −80°C until use. Single-molecule fluorescence in situ hybridization (smFISH) was performed using the RNAscope Multiplex Fluorescent Kit (Advanced Cell Diagnostics, ACD; Hayward, CA) according to the manufacturer’s instructions. The following probes against human transcripts were used: Hs-LEF1-C2 (412991-C2), Hs-TNC-C3 (420771-C3), and Hs-WNT7B (421561). TSA-conjugated Opal 570 and Opal 650 fluorophores were used for signal detection and were interchanged across experimental runs with no differences in outcome. The 488 nm channel was intentionally left unoccupied in smFISH-only experiments due to higher autofluorescence background; it was reserved for concurrent immunofluorescence detection when combined with IF staining.

Prior to hybridization, slides were dried at 60°C for 1 hour and post-fixed in 4% paraformaldehyde (PFA) for 2 hours. Sections were washed three times in PBS (5 minutes each), after which endogenous peroxidase activity was quenched using ACD Hydrogen Peroxide solution for 10 minutes, followed by two washes in distilled water. Target retrieval was performed in 1X Target Retrieval Buffer (ACD) at 95–100°C for 5 minutes. Slides were then washed sequentially in distilled water and 100% ethanol and baked at 60°C for 30 minutes. Sections were rehydrated in PBS for 2 minutes before protease treatment for 15 minutes at 40°C in a HybEZ™ oven.

Probe hybridization and signal amplification were performed according to the manufacturer’s protocol, with one modification: following hybridization, slides were incubated in 5X SSC overnight at room temperature prior to amplification.

Tissue sections were incubated with target probes (2–3 drops per section) for 2 hours at 40°C, followed by two washes in 1X Wash Buffer (ACD) for 2 minutes each. Signal amplification was carried out using the RNAscope Multiplex Fluorescent Detection Kit v2 (ACD, 323110) through sequential AMP-FL reagent cycles; each reagent (∼4 drops per section) was applied for 30 minutes at 40°C in the HybEZ™ oven, followed by two washes in 1X Wash Buffer for 2 minutes at room temperature. Following the final amplification step, slides were washed in PBST, counterstained with DAPI for 30 seconds at room temperature, and coverslipped using Aqua-Mount (Lerner Laboratories). For experiments combining smFISH with immunofluorescence, CD31 primary antibody incubation, secondary antibody incubation, and coverslipping were performed immediately after the in situ hybridization protocol, as described in the “Immunofluorescence Staining” section.

Negative controls, in which fluorophores were applied in the absence of primary probes, were included in all experiments. In postnatal samples, negative controls were additionally used to identify lipofuscin autofluorescence, which was distinguished from genuine RNA signal by its appearance as large, spectrally broad aggregates spanning multiple channels, in contrast to the discrete, diffraction-limited puncta characteristic of single RNA molecules.

### Imaging and quantifications

High-resolution images of hippocampal regions were acquired on a STELLARIS 8 laser scanning confocal microscope with tunable white light laser and hybrid (HyD) detectors, set on a DMI8 inverted stand (Leica Microsystems). Image acquisition was operated with LASX and excitation wavelength and detection bandwidth were optimized and set automatically by the software for each fluorophore. Images were acquired with a 40X oil-immersion Plan Apochromat objective (1.3 NA) in bidirectional scanning mode. For each fluorophore, laser power and detector gain were optimized to maximize signal-to-noise while avoiding pixel saturation. Images were acquired with sequential line scanning to prevent spectral crosstalk. Representative images in figures are displayed as maximum intensity projections. Further adjustments to brightness and contrast were made in Fiji for display. For all quantifications, multiple individual images were sequentially acquired to fully cover the entire region of interest per section. Independent sections from different anterior-posterior levels were analyzed per case and the average number was calculated. Z-stack scan depths of 7-9μm were acquired, with a Z step size of 0.5μm.

#### PROX1 and TBR2 cell densities

PROX1^+^ and TBR2^+^ cells were identified and manually counted by two independent experimenters under blinded conditions. Counts obtained by the two independent experimenters were averaged to obtain a final value for each section. Cell density was calculated by normalizing the number of immunopositive cells to the volume of tissue imaged, derived by multiplying the area of the tissue by the z-stack depth.

#### Volume of NESTIN expression

To quantify the volume occupied by NESTIN expression, the full mediolateral extent of the DG was sampled. The percentage of volume occupied by NESTIN^+^ cells was estimated in confocal z-stacks covering the GCL and GHPZ. First, radial cells were manually selected in each z-stack using the “Threshold” tool (Fiji) to mask only the pixels of the image with NESTIN^+^ staining. Then, the percentage of pixels occupied by NESTIN^+^ filaments was calculated using the Area Fraction parameter of the “Measure” tool (Fiji). Blood vessels were manually selected and excluded from the quantifications using the “Clear” tool (Fiji). The average Area Fraction from 5 (early stages where DG is smaller) to 25 images per z-stack was calculated.

#### Percentage NESTIN/S100B^+^ cells

For quantification of NESTIN^+^/S100B^+^ double-positive cells among radial NESTIN^+^ cells, radial glial cells (RGCs) were identified based on the presence of a single apical process extending toward the molecular layer, with the nucleus visible within the imaging plane. Cells were classified as RGCs when DAPI-labeled nuclei displayed NESTIN expression at the nuclear crown with a filament extending directly from the nucleus. S100B expression was subsequently assessed in identified RGCs. Only cells with nuclei located within the granule cell layer (GCL) or granule cell/hilar proliferative zone (GHPZ) were included in the analysis. NESTIN^+^ filaments without an identifiable nucleus were exluded from the quantification.

GraphPad Prism (v.11.0.2) was used to generate all graphs. Source data for all quantifications is accessible in **Table S2**.

### Bulk RNA Spatial Transcriptomics (GeoMx)

#### Slide preparation and ROI selection

We used nanoString’s GeoMx Digital Spatial Profiler (DSP) to perform spatial transcriptomics at three different developmental stages (early-gestation, mid-gestation and infancy). The whole transcriptome atlas probe panel was used, containing 18,815 single probes for the same number of genes. A GW14 sample was processed and sequenced twice in two separate slides. 1 GW22 sample and two independent GW23 samples were processed in three different slides (grouped as GW22-23), and 1 4-month-old sample was also processed. All steps were performed using RNase-free conditions and DEPC-treated ultrapure water when required. Within two weeks prior to processing the slide on the DSP, 10μm-thick sections were obtained using a cryostat at −20°C, mounted on positively charged Superfrost Plus slides and stored at −80°C until use. Once the slides were taken out from −80°C, they were left at RT for 1-2 minutes and washed with PBS for 5 minutes to remove the excess of OCT. Next, they were baked at 60°C for 1 hour, followed by 1 wash in 50% ethanol, 1 wash in 70% ethanol, and 2 washes in 100% ethanol for 5 minutes each. The slide was left air-drying for 5-10 minutes and target retrieval was performed in target retrieval reagent (10x Invitrogen 00-4956-58) diluted to 1x for 10 minutes at 95°C, followed by a 5-minute wash in PBS. Slides were then incubated in 0.1μg/mL proteinase K (Invitrogen 25530-049) 15 minutes at 37°C and washed in 1x PBS for 5 minutes. Slides were then placed in a humidity chamber humidified with 2X SSC (Dilute from 20x, Sigma-Aldrich, S6639-1L) and incubated with whole transcriptome atlas hybridization probes (nanoString Cat# 121401102) diluted in Buffer R (provided in the GeoMx RNA Slide Prep PCLN kit, # 121300313) in a hybridization oven at 37°C for 16- 20 hours.

Following probe incubation, slides were washed with stringent washes (equal parts formamide and 4x SSC buffer) at 37°C twice for 25 minutes each. Then slides were washed twice in a 2x SSC buffer. Slides were incubated in 200 μL buffer W (provided in the GeoMx RNA Slide Prep PCLN kit, catalog # 121300313) for 30 minutes and incubated in primary antibodies conjugated to specific fluorophores (Anti-Vimentin E-5 conjugated to Alexa Fluor 546, Santa Cruz, #sc-373717 AF546; Anti S100β conjugated to Alexa Fluor 594, Novus, #NBP2-54426AF594; Syto13 nuclear marker, Invitrogen, #S7575) diluted in buffer W (provided in the GeoMx RNA Slide Prep PCLN kit, catalog # 121300313) for 2 hours at RT, to identify specific proteins that served as morphological markers. Slides were washed 2 times in a 2x SSC buffer for 5 minutes and placed in the nanoString GeoMx DSP instrument. Syto13 immunofluorescence was utilized for autofocus of GeoMx imaging and for proper identification of distinct hippocampal structures. Vimentin and S100B were used to locate RGC scaffolds and the fimbria, respectively. ROIs were generated using the polygon tool to discriminate different structures. ROIs in The DVZ, DMS, GCL and fimbria were collected for all samples. ROIs in the GHPZ were collected at GW22-23 and 4 month samples, but not GW14 due to the structure not yet being present at this age. At GW22-23 and 4 months, ROIs collected in the DMS region were named remnant DMS (rDMS) due to the depletion of the stream by mid-gestation. At GW14, a thorough sampling of the CA region was done, covering the distance from ventricle to CA with multiple ROIs. This distance was further subdivided into CA_VZ area for ROIs adjacent to the ventricular zone, CA_SVZ for the region immediately adjacent to the VZ, CAS (CA scaffold) for the region adjacent to CA neuronal layer, and CA for the CA pyramidal neuron layer. Sparse molecular layer regions, CA regions at GW22-23 and ganglionic eminence regions were also collected in the process. Hilar regions that were not adjacent to the GCL were annotated as Hilus, and not considered GHPZ. A total of 282 ROIs were collected across 6 slides. By age, 94 at GW14, 141 at GW22-23 and 47 at 4 months. By region, 10 DVZ, 12 DMS, 50 GHPZ, 78 GCL, 32 fimbria, 5 hilus, 16 molecular layer, 62 CA-related, 14 rDMS, and 3 ganglionic eminence **(Table S3)**. Each ROI, regardless of the surface area covered, contained a minimum of 70 nuclei. Probe identities in each segment were captured via UV illumination and movement to a 96-well plate.

#### Library preparation

Illumina Novaseq 6000 was used for sequencing with a read length of 27 for both reads with reverse sequence orientation in the readout group plate information. Plates were dried down at 95°C and rehydrated in 10 μL nuclease-free water, mixed and incubated at room temperature for 10 minutes. PCR was performed on samples as described in the nanoString GeoMx DSP Readout User Manual using 2 μL PCR master mix, 4 μL primer from the correct wells, and 4 μL resuspended DSP aspirate. KAPA beads (KAPA Pure beads, Roche Cat# 07983298001) were warmed to room temperature for 30 minutes. Libraries were pooled and KAPA beads were added to each pool at a 1.2X ratio to the final pool volume. Two KAPA bead clean ups were performed and pooled libraries were eluted in 0.5 μL elution buffer per collected ROI. Negative control pools were eluted into a 10 μL elution buffer. Pools were diluted 1:8 in elution buffer, and library quality and size were evaluated using the Agilent High Sensitivity DNA chip on the Agilent 2100 Bioanalyzer (Agilent Technologies, Inc.). Sequencing was performed on an Illumina NovaSeq 6000 (SP flow cell) using dual-indexing and paired-end 2 × 27 bp reads, with a 5% PhiX spike-in by volume, generating approximately 400 million reads per lane and a median of 5,508,948 raw reads per ROI. Raw sequencing output was demultiplexed and converted to FASTQ format FASTQ files were subsequently processed into Digital Count Conversion (DCC) files using the GeoMx NGS Pipeline.

#### Data Preprocessing and Quality control

Preprocessing and quality control were performed by adapting the previously reported GeomxTools (v3.5.0) workflow^81^. None of the 282 collected segments failed initial quality control assessment for minimum number of reads (1000), percentage of trimmed, stitched and aligned reads (80%, 80%, and 75%, respectively), minimum sequencing saturation (50%), minimum negative counts (1.01), and minimum area (1000). All ROIs selected had a minimum of 70 nuclei count and no background No Template Control (NTC) counts. After sequencing the data, a total of 18815 were identified in the tissue, including 139 negative control probes. All probes passed initial probe QC, following Nanostring’s standard cutoff criteria. Minimum probe ratio of 0.1 and Grubbs test for outliers. 40 local outliers were identified, all of them corresponding to negative probes. After aggregating the data, resulting in 18677 genes, the limit of quantification (LOQ) was calculated for each gene, setting a cutoff at “2” and a minimum LOQ of 2.

Gene detection above the LOQ threshold was calculated to identify low quality segments and probes. Most segments showed a gene detection ratio (detected/total) above 30%. 31 segments between 15-30% and 2 segments between 5-15%. The two segments that were lower than 15% were excluded from downstream analysis (Both from a 4m sample, one GHPZ and one molecular layer). Only genes present in at least 10% of the segments were selected for downstream analyses (14,483 genes).

To account for systematic variation between segments we performed both Q3 and background normalizations, based on Q3 counts and negative probe counts, respectively. Q3 normalization performed better and was selected for downstream analysis.

#### Data Analysis

GeoMx data were coerced into a Seurat object and the Seurat analysis pipeline (v5.0.3)^82^,originally developed for single-cell RNA sequencing, was adapted for the analysis of this spatial transcriptomics dataset. Briefly, data were normalized using the NormalizeData function and the 2,000 most variable features were identified using the variance stabilizing transformation (VST) method. Data were then scaled and the first 15 principal components (PCs) were calculated and used for downstream analysis. Harmony (v1.2.0)^83^ was applied to correct for batch effects between slides processed on different sequencing scans, prior to neighbor graph construction and cluster identification. Clustering of the full dataset was performed at a resolution of 0.8. GHPZ and GCL ROIs were subsequently subset and independently reanalyzed, repeating variable feature selection, scaling, PCA, Harmony batch correction, and cluster identification at a resolution of 0.6. UMAP was used for visualization throughout.

Differential expression analysis between GHPZ and GCL at mid-gestation, and across developmental stages (early gestation, mid-gestation, and infancy), was performed using the FindMarkers function in Seurat with the MAST statistical test (v1.27.1)^84^, and a minimum expression threshold of 25% of cells per group (min.pct = 0.25). Genes with an adjusted p-value lower than 0.01 and a log2FC higher than 1 were considered significantly upregulated, while genes with a log2FC lower than −1 were considered significantly downregulated.

Gene ontology analysis was performed using the Enrichr web tool^85^ with the GO Biological Process 2025 database. Upregulated genes passing the significance thresholds described above were submitted for each developmental stage (early gestation, mid-gestation, and infancy). GO terms with an adjusted p-value lower than 0.05 were considered significant and visualized using bubble plots. For clarity, significant GO terms were manually grouped into broader biological categories **(Table S4)**. For plotting clarity purposes, the GO terms were grouped in broader categories: Signaling pathway-related (General, Semaphorin-Plexin, growth factor & RTK, WNT, Notch, MAPK, Nitric Oxide (NO), Interleukin 3 (IL3), Cytokine, and ERK signaling), Neuron Maturation, Synaptic Maturation, Nervous System Development, General Development, Cell Adhesion, Cell Proliferation, Cell Migration, Cell Differentiation, Cell Metabolism, Immune Response, Extracellular Matrix.

### Single-Cell RNA Spatial Transcriptomics (CosMx SMI)

#### Slide preparation

Two independent 10 µm-thick cryosections from a GW23 human brain sample were post-fixed in 4% PFA, mounted onto positively charged Superfrost Plus slides, and stored at −80°C until use. Slides were processed following the Fresh Frozen section of the CosMx SMI Slide Preparation manual (Bruker Spatial Biology, MAN-10184-04) using the CosMx FFPE Slide Prep Kit (RNA) (121500006). Briefly, slides were washed thrice in PBS and baked at 60°C for 30 minutes to enhance tissue adhesion. To rehydrate, slides were washed in PBS, followed by 50% ethanol for 5 minutes and then stored in 70% ethanol at 4°C overnight. The following day, slides were washed in 100% ethanol twice and then left on a benchtop to air dry for 10 minutes. Antigen retrieval was performed in 1X NanoString Target Retrieval Buffer at 100°C for 10 minutes using a pressure cooker, followed by sequential washes in DEPC water and 100% ethanol for 3 minutes. Following a 30-minute air dry step, tissue permeabilization was achieved by following the manual’s “Fresh Frozen tissue types” guide, which entailed using a fresh dilution of Proteinase K at 3 μg/mL working concentration. Once prepared, 400μL of this digestion solution was directly applied onto the tissue, and samples were incubated for 30 minutes at room temperature. Following two PBS washes, freshly prepared and vortexed fiducials were applied to the tissue sections for 5 minutes, and tissue sections were protected from light from this point forward. After another PBS wash, a quick fixation was done by keeping the slides in 10% neutral buffered formalin solution (Sigma-Aldrich, HT5012) for 1 minute. The reaction was quenched by washing with freshly prepared Tris-Glycine buffer twice, for 5 minutes each. Another PBS wash was performed, and tissue sections were incubated with 100mM NHS-Acetate mixture for 15 minutes at room temperature to quench reactive amine groups. After a 2X saline-sodium citrate (SSC) wash, the CosMx 6k Discovery Panel (Nanostring, 121500041) was applied. This pool of recently denatured and crash cooled in-situ hybridization probes targeting a panel of 6175 human gene transcripts (+20 non targeting controls) was added along with RNase Inhibitor, Buffer R, and rRNA segmentation markers, and incubated overnight at 37°C. The following day, two stringent washes with 50% formamide were performed in a 37°C water bath for 25 minutes each, and slides were washed in 2X SSC twice. Tissue sections were incubated with a DAPI nuclear stain for 15 minutes, washed in PBS thrice, and then incubated with neural-specific cell segmentation markers (GFAP and Histone) for 1 hour. Three more PBS washes were done, and samples were incubated with 100mM NHS-Acetate mixture for 15 minutes at room temperature again. Slides were washed with 2X SSC and flow cells were applied before CosMx machine loading. For the flow cell configuration setup, “Configuration B: 90 seconds” was used for the pre-bleaching profile, and “Configuration B: Human Neural Tissue” was used for the cell segmentation profile. Following the initial scan, 117 square fields of view (FOVs) were placed over the hippocampus regions of interest and the run was initiated as described in the CosMx® SMI Instrument manual (Bruker Spatial Biology, MAN-10161-08).

#### Data analysis

Following scan completion and upload to NanoString’s cloud AtoMx Spatial Analysis Platform, “studies” were created for each slide and the following flat csv files were exported for in-house analysis: count matrix, cell metadata, transcripts, polygons, and FOV positions. Scanpy^86^ was used for all downstream preprocessing, dimensionality reduction, clustering, analysis, and visualization. Cells with fewer than 300 total transcripts were excluded from the dataset (∼13% of cells filtered out). Following all pre-processing, slide #1 retained 38,306 cells and slide #2 retained 23,809 cells, and both slides contained ∼1,800 transcripts per cell on average (mean). Counts were normalized to 10,000 per cell with ‘sc.pp.normalize_total’ and log-transformed with ‘sc.pp.log1p’, and highly variable genes, PCA, the neighborhood graph, the UMAP embedding, and Leiden clustering were then computed with Scanpy default parameters (‘sc.tl.pcà, ‘sc.pp.neighbors’, ‘sc.tl.umap’, ‘sc.tl.leiden’). Initial Leiden clusters were annotated to broad cell types by inspection of canonical marker genes. One Leiden cluster lying near the center of the UMAP with diffuse expression characteristic of low-quality cells was removed and the embedding was recomputed on the remaining cells. This initial annotation was then refined through several rounds of cluster-restricted subclustering on the existing PCA representation: the RGC cluster was sub-clustered (‘sc.tl.leiden(resolution=0.15)’) and the EOMES-high subcluster was identified as Intermediate Neural Progenitors (INPs); inhibitory neurons were re-examined and subdivided with the canonical MGE markers (LHX6, SST, NKX2-1) versus CGE markers (NR2F2, PROX1, CALB2), and an intermediate “CA neuron” cluster was sub-clustered into two Leiden groups and reassigned as CA1 Immature Neurons and CA3 Immature Neurons. This iterative refinement produced the final 21 cell-type annotation used for slide #1. The same procedure was applied independently to a replicate GW23 slide #2 for the supplementary figure. To validate these annotations in a complementary reference frame, we integrated the CosMx data with our age-matched (GW21 + GW23) snRNA-seq samples using Harmony^83^: the two modalities were scaled separately, concatenated, and embedded via PCA followed by ‘harmony_integratè on the batch covariate (‘max_iter_harmony=20’); a joint neighborhood graph was then computed on the Harmony-corrected embedding (‘n_pcs=20’, ‘n_neighbors=20’) and visualized with UMAP. snRNA-seq cell-type labels were projected onto the CosMx cells in this integrated space using a distance-weighted k-nearest-neighbors classifier (k=20) with a minimum class-probability of 0.5 required for assignment. Marker gene expression and clear anatomical patterns in single-cell spatial distribution were used for accepting cell type assignments **(Table S5)**.

To map proliferative activity onto the hippocampus at a sparse single-cell level, each CosMx cell was classified as G1, S, or G2/M by a two-tier marker-based procedure: cells passing a “cycling” gate on a panel of proliferation markers (MKI67, TOP2A, CDK1, UBE2C) or on a panel of S-phase replication markers (PCNA, MCM2/4/5/6, TYMS, GMNN, and others) were considered cycling. Next, the S- versus G2/M-specific marker modules (CENPF, NUSAP1, BIRC5, TPX2, the AURKA/B–CCNB2–KIF11 mitotic block, etc.) were z-scored across these cycling cells and compared per individual cell, with the dominant module determining phase. Cells failing these gates were assigned to G1, and phase labels were projected onto the tissue. To compare cell populations between regions within the tissue, we performed an inside-vs-outside differential gene expression contrast for two cell types in the GW23 section (Figure 3G, 3H). Anatomical regions were defined by hand-selected polygons on the spatial (‘CenterX_global_px’, ‘CenterY_global_px’) coordinate system and isolated with the ‘matplotlib.path.Path.contains_points’ function. For RGCs, two polygons were drawn: one enclosing the DG interior and one enclosing the adjacent ventricular zone immediately outside the dentate. The RGC cells lying inside each polygon were used as the two comparison groups. For Proliferating cells, a single dentate polygon was drawn and Proliferating cells inside it were compared against all remaining Proliferating cells distributed elsewhere in the tissue section. Differentially expressed genes were called by Wilcoxon rank-sum test (‘sc.tl.rank_genes_groups’, ‘method=’wilcoxon’’, ‘pts=Truè). Genes were considered differentially expressed at |log₂ fold change| ≥ 0.5, fraction of cells expressing in the in-group ≥ 0.10, and Benjamini–Hochberg-adjusted p < 0.05.

### Single Nucleus RNA sequencing (10x genomics)

#### Nuclei isolation and library preparation

Prior nuclei isolation and single nucleus RNA sequencing, RNA quality was assessed by extracting total RNA from each case and measuring RNA Integrity Number (RIN) using the Agilent 2100 Bioanalyzer (Agilent Technologies, Inc.). Only samples with RIN ≥ 7 were included in the study.

For the first batch of samples (GW15, GW18, GW19, GW21, 2weeks, 4months, 7 months and 38 years) nuclei isolation was performed adapting a protocol described before^87^. Cryosectioned brain tissue (30-40mg) was homogenized in 5 mL RNAase-free lysis buffer (0.32M sucrose, 5 mM CaCl2, 3 mM MgAc2, 0.1 mM EDTA, 10 mM Tris-HCl, 1 mM DTT, 0.1% Triton X-100 in DEPC-treated water) using a glass Dounce homogenizer (Thomas Scientific, Cat # 3431D76) on ice. The homogenate was loaded into a 30 mL thick polycarbonate ultracentrifuge tube (Beckman Coulter, Cat # 355631) and 9 mL of sucrose solution (1.8 M sucrose, 3 mM MgAc2, 1 mM DTT, 10 mM Tris-HCl in DEPC-treated water) was added to the bottom of the tube. The sample was centrifuged at 107,000 g for 2.5 hours at 4°C. After aspirating the supernatant, the pellet was incubated in 250 µL DEPC-treated PBS for 20 minutes on ice before resuspension. The nuclear suspension was filtered twice through a 30 µm cell strainer, and nuclei were counted using a hemocytometer and diluted to 2,000 nuclei/µL. Single-nucleus capture was performed on the 10x Genomics Single-Cell 3’v3 system, following the manufacturer’s protocol. Libraries were pooled and sequenced on a NovaSeq 6000 S2 flow cell (average depth 40,000 reads/nucleus). Raw sequencing data were processed using Cell Ranger v[3.0.2] aligned to the [hg38/mm10] reference genome.

The second batch (GW18, GW23, 2weeks and 7months) was prepared following the 10x Genomics demonstrated protocol (CG000365). Cryosectioned brain tissue (30-40mg) was homogenized in 1 mL NP40 Lysis buffer (10 mM Tris-HCl (pH 7.4), 10 mM NaCl, 3 mM MgCl₂, 0.1% NP40, 1 mM DTT (Millipore Sigma, 646563), and RNase inhibitor) using a glass Dounce homogenizer (Millipore-Sigma, DWK885300-0001). The sample was kept on ice throughout the procedure. After a 5-minute incubation, the homogenate was filtered through a 70 µM Flowmi cell-strainer (SP-Bel Art, H13680-0070) into a 50mL conical tube, then transferred into a 1.5mL tube. After centrifugation (500 g for 5 minutes at 4°C), the supernatant was discarded, and the pellet was incubated for 5 minutes in 500 µl permeabilizing solution (PBS+1%BSA+1U RNase inhibitor (Millipore-Sigma, 3335399001)). After resuspension, the sample was centrifuged again (500 g for 5 minutes at 4°C), and the supernatant was discarded. The pellet was resuspended in 1mL permeabilizing solution, followed by the addition of 10ul 7AAD (Sigma-Aldrich, SML1633) to identify nuclei. Fluorescence-activated cell sorting (FACS) was then performed to enrich nuclei and exclude whole-cells. A 100 µL aliquot of the sample was retained as a negative control during FACS. Nuclei were collected into 15 mL tubes containing 500 µL of permeabilizing solution, then centrifuged (500 g for 5 minutes at 4°C) and resuspended in 1X Lysis Buffer (10 mM Tris-HCl (pH 7.4), 10 mM NaCl, 3 mM MgCl₂, 0.1% NP40, 0.1% Tween-20, 0.01% digitonin (Thermofisher, BN2006), 1% BSA, 1 mM DTT, and RNase inhibitor) for gestational samples (GW18 and GW23) or 0.1X Lysis Buffer for postnatal samples (2weeks and 7months). After a 2 minute incubation on ice, 1mL wash buffer (10 mM Tris-HCl (pH 7.4), 10 mM NaCl, 3 mM MgCl₂, 0.1% Tween-20, 1% BSA, 1 mM DTT, and RNase inhibitor) was added, and the suspension was pipetted five times before centrifugation (500 g for 5 minutes at 4°C). Nuclei were counted using the Countess II automated cell counter (Thermo Fisher Scientific, AMQAX1000) and diluted to 4,000 nuclei/uL in Diluted Nuclei Buffer (1x Nuclei Buffer (10x Genomics, Chromium Single Cell Multiome ATAC + Gene Expression kit), 1mM DTT, 1U/ul RNAse inhibitor, Nuclease-free water (Fisher, AM9937). Nuclei quality was assessed by DAPI staining of a sample aliquot (20-50 µL) and fluorescence microscopy using 40X and 63X objectives. The preservation of membrane integrity and lack of blebbing were used to determine healthy nuclei. Single-nucleus capture and library preparation followed the 10x Genomics Chromium Next GEM Single-Cell Multiome ATAC+Gene Expression User Guide (#CG000338). Libraries were pooled and sequenced on a NovaSeq 6000 S4 flow cell (average depth 40,000 reads/nucleus). Raw sequencing data were processed using Cell Ranger ARC aligned to the [hg38/mm10] reference genome.

#### Data processing and filtering

Data from each sample were filtered based on read count, gene expression, and mitochondrial and ribosomal content. Violin plots for each of the variables were run before and after performing unbiased clustering using the Seurat pipeline^82^. Scatter plots were also run for side-by-side comparisons between variables. Nuclei with low read counts, low gene expression, or high mitochondrial/ribosomal content were excluded. For gestational samples in the first batch, nuclei expressing fewer than 500 or more than 7,000 (GW15), 6,000 (GW19) or 7,500 (GW21) genes were excluded. For GW18, only one subcluster (Cluster 9) was retained because it exhibited meaningful differential gene expression and spatial differentiation from the other nuclei, due to a low number of high-quality nuclei. For postnatal samples, the gene expression limits were set to < 500 and > 10,000 genes. Nuclei with greater than 10% ribosomal content were excluded from GW15 and GW19, nuclei with more than 2.5% mitochondrial content were excluded from samples 2weeks, 4months, 7months, and 38years, and those with more than 1% mitochondrial content were excluded from GW21. The nuclei with higher mitochondrial/ribosomal content than the chosen threshold generally exhibited the lowest gene counts. The second batch was filtered excluding nuclei with fewer than 500 genes, more than 7,000 genes, or greater than 40% ribosomal content on each sample.

After filtering, individual matrices were combined prior to pre-processing and clustering.

#### Data integration and annotation

Our base embedding was created in the scanpy^86^ package, by running PCA and then using harmony^83^ to correct batch in the PCA space between our two primary sample sequencing events (2019 and 2022). K-nearest neighbors (knn) was then run using the top 20 PCs sorted by variance and k=20 neighbors. The knn graph was then used to compute a two-dimensional UMAP embedding. Cell type identities were assigned using a combination of unsupervised clustering and canonical marker gene expression. Clustering was performed across multiple resolutions, and individual clusters were subset and re-clustered where necessary to resolve closely related subtypes and confirm identity. Once a biologically meaningful annotation granularity was achieved, fine-level cell types were grouped into broader annotation categories for downstream analyses. Annotations were generated at three hierarchical levels. Level 1 annotations contained major cell classes including NPCs, macroglia, excitatory and inhibitory neurons, and a category with distinct cell populations classified as others. Level 2 annotations represented broad cellular classes within the Level 1 annotations, including RGCs, INPs, and proliferating cells within NPCs, astrocytes, GPCs, oligodendrocytes, and OPCs within macroglia, CA, GC, and a population broadly categorized as immature neurons within excitatory neurons, CGE- and MGE-derived neurons within inhibitory neurons, and microglia, endothelial cells, ependymal cells, choroid plexus cells, VLMCs, a population of neurons showing thalamic signature possibly derived from adjacent thalamic sectioning, and a small subset of unidentified cells within others. Level 3 annotations represented finer transcriptional subtypes within these broader classes, derived from additional subsetting and reclustering of each of the L2 categories. Annotation details can be observed in **Table S6**.

#### RGC only pseudotime

The data was subset to only the RGC subtypes (RGC1, RGC2, RGC2-N, RGC2-A, and RGC3) and the neighbors graph was recomputed on the harmony corrected PCA space using n-neighbors=20 and n pcs = 20. Pseudotime was recomputed from a root not in RGC1 from the GW15 donor. Glial cell types (RGCs_1, RGCs_2, RGCs_2-A, RGCs_3) were again inverted for bidirectional pseudotime) with glial extending to −1 and neuronal to 1, as previously described^88^.

#### NPCs, astrocyte and oligodendrocyte lineages and excitatory neurons pseudotime

This analysis consisted of RGCs, proliferating cells, GPCs, astrocytes, OPCs, oligodendrocytes, INPs, immature neurons, granule neurons of the DG, and CA region neurons of the hippocampus. A considerable batch was observed between GW15 and all other ages so to preserve developmental trajectory across ages batch was recorrected by harmony^83^ using 3 batches (2019 dataset, 2022 dataset, and GW15) A root node was then set in a RGC1 cell from GW15 where diffusion based pseudotime was then computed using the Scanpy implementation^86^. Glial cell types (RGCs_1, RGCs_2, RGCs_2-A, RGCs_3, GPC, Astrocytes, OPC, and ODC) pseudotime values were again inverted to create a bidirectional pseudotime where glial developmental time moves towards −1 and neuronal towards 1.

#### Signaling pathway enrichment in RGCs

To identify cell-type-enriched signaling pathways, differential gene expression analysis was performed for each of the five neuronal subtypes using a one- versus-all comparison (t-test implementation in Scanpy). Genes with an adjusted p-value < 0.01 and Log2FC > 0.5 were considered significantly upregulated. The resulting upregulated gene lists were submitted to Enrichr web tool^85^ for pathway enrichment analysis using the KEGG 2024 and Reactome 2024 databases. Pathways with an adjusted p-value < 0.05 were considered significantly enriched **(Table S7)**.

#### Mouse-Human Integration

Integrating our dataset with a developmental mouse dataset from the same brain region^52^, first involved mapping mouse genes to their human orthologs and retaining genes with one-to-one orthologous relationships. Highly variable genes (HVGs) from both datasets were then calculated using scanpy’s highly variable genes function, and both datasets were subsequently restricted to the intersection of both HVG sets. Harmony was then run on 3 batches (2019, 2022, and Hochgerner et al. 2018) to correct the batch between species as well as retaining the batch correction between the two sequencing runs within our dataset. This integrated dataset was then reduced to the same cell types observed in the above “NPCs, astrocyte and oligodendrocyte lineages and excitatory neurons pseudotime” analysis.

#### DecoupleR

The python decoupleR package^51^ was run on the RGC subset using decoupler.op.collectri with organism=human and decoupler.mt.ulm with tmin parameter set to 5.

### Human adult and early postnatal dataset (Dumitru et al., 2025) integration

#### Dataset stratification and preprocessing

Single-cell RNA sequencing data generated in this study were integrated with adult hippocampal single-cell transcriptomic data from Dumitru et al. (2025)^24^ to investigate developmental and adult hippocampal cell populations. Donor samples from Dumitru et al. were split into early postnatal and adult groups based on donor age. Donors aged 0–5 years were classified as early postnatal, and donors ranged from 13 to 78 years of age were classified as adults.

To maximize retention of rare developmental populations, no additional cell-level quality-control filtering was applied beyond the filtering performed in the original datasets. Following dataset merging and preliminary clustering, donor DG-450 was identified as a marked outlier. Manual annotation revealed that 93.5% of cells from this donor were classified as immature neurons, while several major hippocampal cell populations, including astrocytes, MGE-derived interneurons, microglia, oligodendrocytes, and OPCs, were completely absent. Because this cellular composition differed substantially from all other samples, DG-450 was excluded from subsequent analyses. Data preprocessing was performed using Seurat v5.4.0^82^. Following the standard workflow, expression matrices were log-normalized using NormalizeData, highly variable genes were identified using FindVariableFeatures, expression values were scaled using ScaleData, and principal component analysis (PCA) was performed using RunPCA.

#### Integration of gestational and postnatal datasets

To generate a developmental reference atlas, gestational and postnatal samples were combined into a single object. Batch correction and integration were performed using Harmony v1.2.4^83^ on the first 30 principal components. Individual donor identity was used as the integration variable to reduce donor-specific effects while preserving biological variation.

The Harmony embeddings were used to construct a shared nearest-neighbor graph using FindNeighbors, followed by unsupervised clustering using FindClusters. A two-dimensional UMAP embedding was generated from the Harmony-corrected space using the first 30 Harmony dimensions with RunUMAP. The UMAP model was retained to allow projection of external datasets into the developmental reference space.

#### Projection of adult cells into the developmental reference

Adult samples were analyzed separately from the gestational and postnatal reference. Adult cells underwent the same preprocessing workflow, including normalization, variable feature selection, scaling, and PCA.

To compare adult populations directly with developmental populations, adult cells were projected into the gestational/postnatal reference using Seurat’s reference mapping framework. Transfer anchors were identified using the FindTransferAnchors function with PCA projection using the first 30 principal components.

Adult cells were subsequently mapped onto the developmental reference using MapQuery. The Harmony-derived UMAP model generated from the reference was used to embed adult cells into the same low-dimensional space. Cluster assignments were transferred to adult cells, enabling direct comparison of developmental and adult populations within a common transcriptional framework.

#### Cell type annotation

Initial annotations were derived from cluster identities within the developmental reference and subsequently refined through examination of established marker genes previously described. Because the objective of this study was to characterize developmental and adult cell populations at higher resolution, all downstream analyses were performed using the Level 3 annotation scheme that included RGC subtypes and neuronal populations divided by immature and mature states.

#### Cell type composition analysis

To compare cell-type composition between early postnatal and adult hippocampal samples, analyses were restricted to cells originating from the Dumitru et al. dataset. Samples generated in this study were excluded from composition analyses to minimize confounding from differences in dataset origin and developmental stage.

For each annotation category, the proportion of total cells assigned to each cell type was calculated separately for early postnatal and adult samples. Cell-type composition was evaluated. Results are presented as descriptive percentages of cells within each age group. Canonical marker gene expression was evaluated to validate cell type annotations and characterize transcriptomic differences between early postnatal and adult populations. Gene expression patterns were visualized using violin plots.

## Supplemental Figures and Tables

**Figure S1.** Extended cytoarchitectural characterization of the human hippocampus from early- to mid- gestation.

**Figure S2.** Extended spatial transcriptomic sampling and profiling, with complementary cellular and ultrastructural characterization of the DG. Related to Figure 2.

**Figure S3.** Additional data supporting single-cell spatial transcriptomic characterization of the human hippocampus at mid-gestation. Related to Figure 3.

**Figure S4.** Extended transcriptomic analysis of cell populations in the human developing hippocampus. Related to Figure 4.

**Figure S5.** Integration of snRNAseq dataset with a published postnatal dataset and extended characterization of RGCs in the developing GCL/GHPZ. Related to Figure 5.

**Figure S6.** Extended analyses of RGC subtype gene programs, conserved trajectories, and spatial expression. Related to Figure 6.

**Table S1.** Human Brain Specimens used in this study.

**Table S2.** Cellular quantification data.

**Table S3.** Metadata of the bulk-RNA spatial transcriptomics (GeoMx) dataset..

**Table S4**. Gene Ontology enrichment analysis (Biological process) of differentially expressed genes in GCL and GHPZ across developmental stages, related to Figure 2.

**Table S5.** Metadata of the single-cell RNA spatial transcriptomics (CosMx) dataset.

**Table S6.** Metadata of the snRNAseq dataset

**Table S7.** Gene ontology enrichment analysis of differentially expressed genes between RGC types using KEGG and Reactome databases, related to Figure 6.

**Figure S1. Extended cytoarchitectural characterization of the human hippocampus from early- to mid- gestation. Related to** Figure 1**. (A)** Immunofluorescence for NESTIN and S100B in the GW14 dorsal (left), GW14 ventral (center) and GW18 (right) HPC. **(B)** Left: Immunofluorescence for NESTIN and S100B in the DVZ at GW18. Right: TEM image of the DVZ at GW17. Yellow outlined inset highlights a leading process oriented perpendicular to the ventricular wall **(C)** Immunofluorescence for TBR2 in the GW14 dorsal HPC and GW18 HPC. **(D)** Left: immunofluorescence for Ki67 in the GW14 ventral HPC. Right: Ultrathin section of the GW17 HPC with red dots marking identified mitotic cells and TEM micrographs of mitotic cells (pseudo-colored in blue) containing intermediate filaments (arrows) and glycogen granules (arrowheads) in their cytoplasm **(E)** Immunofluorescence for PROX1 and DCX in the GW14 dorsal HPC. **(F)** Immunofluorescence for NESTIN and S100B in the GW18 DG. **(G)** Immunofluorescence for HOPX and GFAP in the GW14 ventral HPC. **(H)** immunofluorescence for NESTIN and S100B in the GW22 (left) and GW30 (right) HPC. **(I)** Top: TEM images showing a uniciliated cell in the DVZ at GW17 left) and a multiciliated cell in the DVZ at GW21 (right). Bottom: Quantification of uniciliated and multiciliated in the DVZ at GW17, GW19 and GW21. **(J)** TEM image of the DVZ at GW22. **(K)** Immunofluorescence for NESTIN and S100B in the DVZ at GW22 **(L)** Immunofluorescence for GAD1 and DCX in the ventricular wall and CA1 region at GW30. Yellow box highlights a cell co-expressing GAD1 and DCX in CA1. **(M-N)** Immunofluorescence for TBR2 (M) and Ki67 (N) in the GW22 and GW30 HPC, respectively. **(O)** Immunofluorescence for HOPX and GFAP in the DG at GW30 (top) with high magnification of the molecular layer. Yellow arrows show cells co-expressing HOPX and GFAP with astrocyte morphology (bottom) <u>Abbreviations:</u> CA, cornu ammonis; D, dorsal; DG, dentate gyrus; DMS, dentate migratory stream; DVZ, dentate ventricular zone; Fi, fimbria; GCL, granule cell layer; GHPZ, granular-hilar progenitor zone; GW, gestational week; HPC, hippocampus; L, lateral; VZ, ventricular zone. <u>Scale bars:</u> (A, C, D, E, G, H, L, M, N) 300 µm; (B, F, J, K, O) 100 µm; (J and O Close ups) 20 µm; (D, TEM) 2 µm (mitosis) and 500nm (Close up).

**Figure S2. Extended spatial transcriptomic sampling and profiling, with complementary cellular and ultrastructural characterization of the DG. Related to** Figure 2**. (A)** Immunofluorescence for the morphological markers (Vimentin, S100B) and nuclear marker SYTO3 used for anatomical and cellular landmark identification in the human hippocampal cases processed through the GeoMx Digital Spatial Profiler workflow. **(B)** Top 10 genes defining unsupervised clusters of the 280 ROIs collected through the GeoMx workflow. **(C)** Heatmap of the top downregulated genes in combined GCL and GHPZ ROIs at GW14, GW22-23, and 4 months. Selected neurodevelopmentally relevant genes are highlighted. **(D)** Ridgplots showing expression of selected genes by age. **(E)** Gene Ontology biological process enrichment analysis of genes downregulated in combined GCL and GHPZ ROIs at GW14, GW22-23, and 4 months. **(F)** Left: Electron microscopy images of the GCL in a neonatal case. Right: Examples of synapses occurring at the nuclear level in the neonatal GCL. **(G)** Immunofluorescence for NESTIN showing the medial GCL and GHPZ at GW34, GW41, 2 years and 7 years. The yellow dashed line outlines the GCL. <u>Abbreviations:</u> CA, cornu ammonis; D, dorsal; DG, dentate gyrus; DMS, dentate migratory stream; DVZ, dentate ventricular zone; Fi, fimbria; GCL, granule cell layer; GHPZ, granular-hilar progenitor zone; GW, gestational week; L, lateral; V, ventral. <u>Scale bars:</u> (F) 10 µm GCL image, 1µm nucleus and 200nm high-magnification. (G) 100 µm.

**Figure S3. Additional data supporting single-cell spatial transcriptomic characterization of the human hippocampus at mid-gestation. Related to** Figure 3**. (A)** Immunofluorescence for GFAP, Histone3 and ribosomal RNA in one example GW23 slide used in the CosMx spatial transcriptomics workflow. **(B)** Spatial mapping (left) and UMAP representation (right) of 19 individual cell types identified by unsupervised leiden clustering and manual annotation. **(C)** Dotplot highlighting the expression of canonical markers across the 21 clusters identified in B. **(D)** Spatial mapping of cells classified as INPs and the INP marker EOMES. **(E)** Left: Spatial mapping of cells classified as proliferating cells. Right: Cell cycle score analysis and spatial mapping of cells classified as G1, S or G2M phase cells. **(F)** Spatial mapping of cells classified as RGCs and the RGC markers NESTIN and TNC. **(G–H)** Left: Spatial mapping of proliferating cells (G) and radial glia-like cells (H) colored by location inside (red) or outside (blue) the DG. Right: Corresponding volcano plots comparing cells from inside versus outside the DG; genes with adjusted p < 0.05 and |log₂ fold change| > 0.5 were considered differentially expressed. <u>Abbreviations</u>: CA, cornu ammonis; CGE, caudal ganglionic eminence; fi, fimbria; GCL, granule cell layer; GE, ganglionic eminence; GHPZ, granular-hilar progenitor zone; GPC, glial progenitor cell; GW, gestational week; HPC, hippocampus; INP, intermediate neural progenitor; MGE, medial ganglionic eminence; RGC, radial glia cell; UMAP, uniform manifold approximation and projection; VZ, ventricular zone.

**Figure S4. Extended transcriptomic analysis of cell populations in the human developing hippocampus. Related to** Figure 4**. (A-C)** UMAP embeddings of 28,122 nuclei across nine hippocampi annotated by sample age (A), superclass (B), and astrocytic andRGC genes of interest (C). **(D)** Expanded dotplot highlighting the expression of canonical markers across the 20 cell types identified in A. **(E-F)** Left: UMAP embedding of nuclei classified as CA neurons (E) and inhibitory neurons (F), annotated by cell subclass and developmental stage. Right: Expression of canonical markers defining each population, including immature (DCX, STMN1, SOX4) and mature (RBFOX3) neuronal markers. **(G)** Left: UMAP embedding of integrated spatial transcriptomic data (replicate GW23 slide) and snRNA-seq datasets (GW21 and GW23). Right: Projection of snRNA-seq–defined cell types onto the integrated spatial–snRNA-seq embedding. (G) Spatial projection of snRNA-seq–defined cell types in the spatial context, with selected populations highlighted. <u>Abbreviations</u>: CA, cornu ammonis; CGE, caudal ganglionic eminence; fi, fimbria; GCL, granule cell layer; GHPZ, granular-hilar progenitor zone; GPC, glial progenitor cell; GW, gestational week; INP, intermediate neural progenitor; MGE, medial ganglionic eminence; RGC, radial glia cell; UMAP, uniform manifold approximation and projection; VZ, ventricular zone.

**Figure S5. Integration of snRNAseq dataset with a published postnatal dataset and extended characterization of RGCs in the developing GCL/GHPZ. Related to** Figure 5**. (A)** Top: UMAP embedding of the integration of our snRNA-seq data with the Dumitru et al. (2025) infant and childhood dataset (left), and projection of Dumitru et al. (2025) adult samples onto the integrated UMAP (right). Bottom: Bar graph showing percentage of cells per cell type within the integrated infant/childhood dataset (left) and adult projected dataset (right). **(B)** Violin plots showing canonical marker genes for selected cell types across both the integrated infant/childhood dataset (left) and adult projected dataset (right). **(C)** Immunofluorescence for NESTIN and S100B in the GCL and GHPZ region at GW34, 2 years and 7 years. **(D)** Immunofluorescence for NESTIN, S100B and VIMENTIN in the GCL/GHPZ at GW37. <u>Abbreviations</u>: CA, cornu ammonis; CGE, caudal ganglionic eminence; GCL, granule cell layer; GHPZ, granular-hilar progenitor zone; GPC, glial progenitor cell; GW, gestational week; INP, intermediate neural progenitor; MGE, medial ganglionic eminence; RGC, radial glia cell; UMAP, uniform manifold approximation and projection; VZ, ventricular zone. <u>Scale bars:</u> (C, D) 100 µm; (D) 20 µm for High magnification images.

**Figure S6. Extended analyses of RGC subtype gene programs, conserved trajectories, and spatial expression. Related to** Figure 6**. (A)** Dotplot of genes driving signaling pathway enrichment of WNT (RGC2A and RGC3), Hippo (RGC2A) and Notch (RGC3) in Figure 6C. **(B)** Heatmap illustrating top 30 genes with enriched inferred activity in DecoupleR analysis per RGC subtype **(C)** Top: UMAP embedding of the integrated mouse dataset from Hochgerner et al. and our human snRNA-seq dataset, colored by cell type and species. Bottom: UMAP embeddings of the integrated mouse–human dataset highlighting sample age, shown separately for the human and mouse cells. **(D)** Top: UMAP embeddings of integrated RGCs, GPCs and astrocytes colored by cell type, dataset of origin, and pseudotime trajectory. Bottom: Genes with the strongest negative and positive associations with pseudotime, representing the major transcriptional programs driving progression across the integrated RGC–GPC–astrocyte trajectory. **(E)** Combined smFISH and immunohistochemistry for LEF1, WNT7B (RNA) and CD31 (protein) in the GCL and GHPZ regions at GW18, GW37, 3weeks and 9 years. The square box in the 9 year picture shows a high magnification image of a CD31-positive blood vessel expressing LEF1 (orange arrow). Red arrows signal lipofuscin aggregates. <u>Abbreviations:</u> CA, cornu ammonis; D, dorsal; FA, Force atlas; GC, granule cell; GCL, granule cell layer; GHPZ, granular-hilar progenitor zone; GPC, glia progenitor cell; GW, gestational week; INP, intermediate neural progenitor; L, lateral; NB, neuroblast; nIPC, neural intermediate progenitor cell; RGC, radial glia cell; RGL, radial glia like; RGLy, Radial glia like young; UMAP, uniform manifold approximation and projection. <u>Scale bars:</u> (E) 50 µm and 20 µm for high magnification images.

