## Supplementary figures and images for "Spatial and Molecular Progression of Neural Progenitor Cells in the Developing Human Dentate Gyrus"

### Supp Info

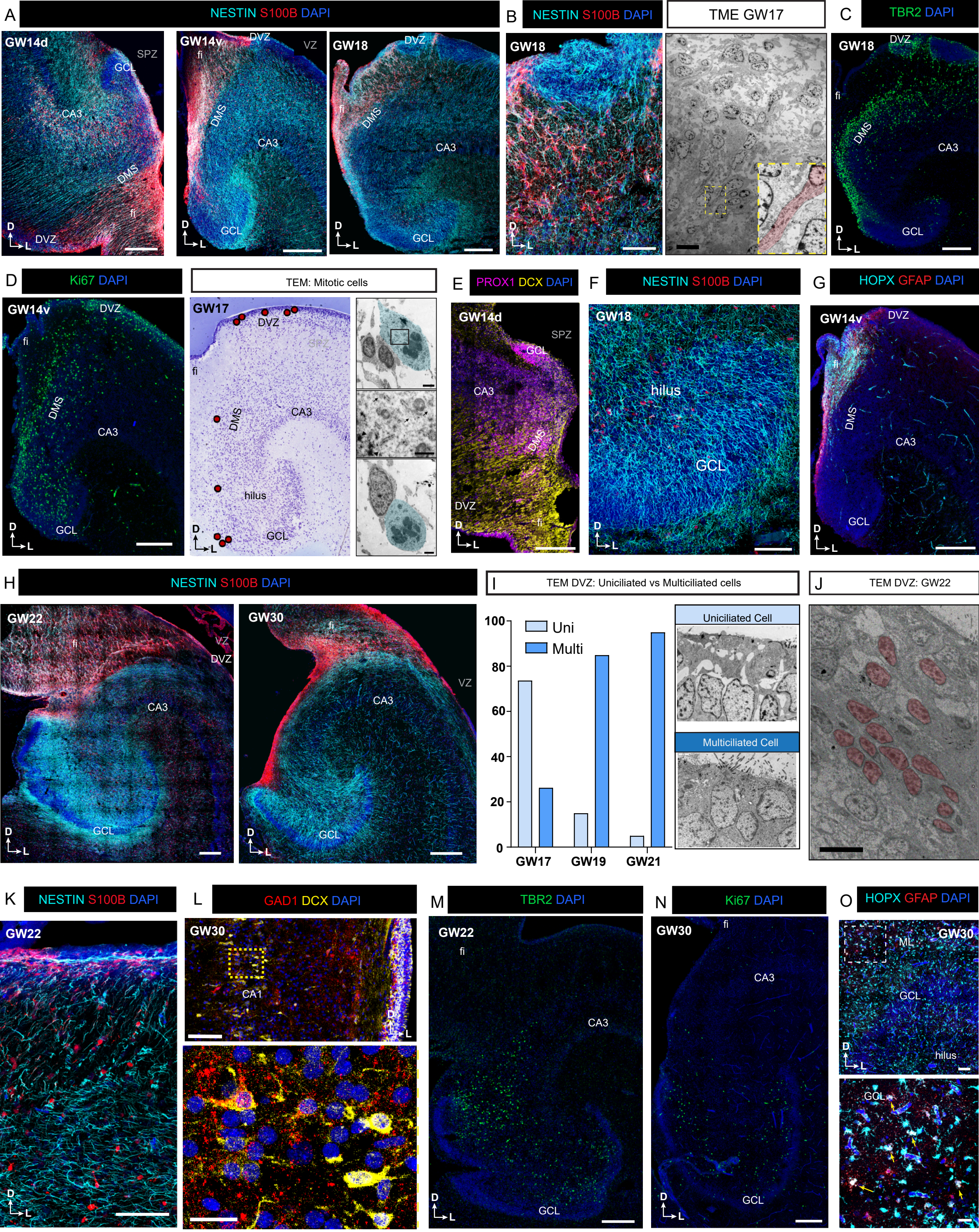

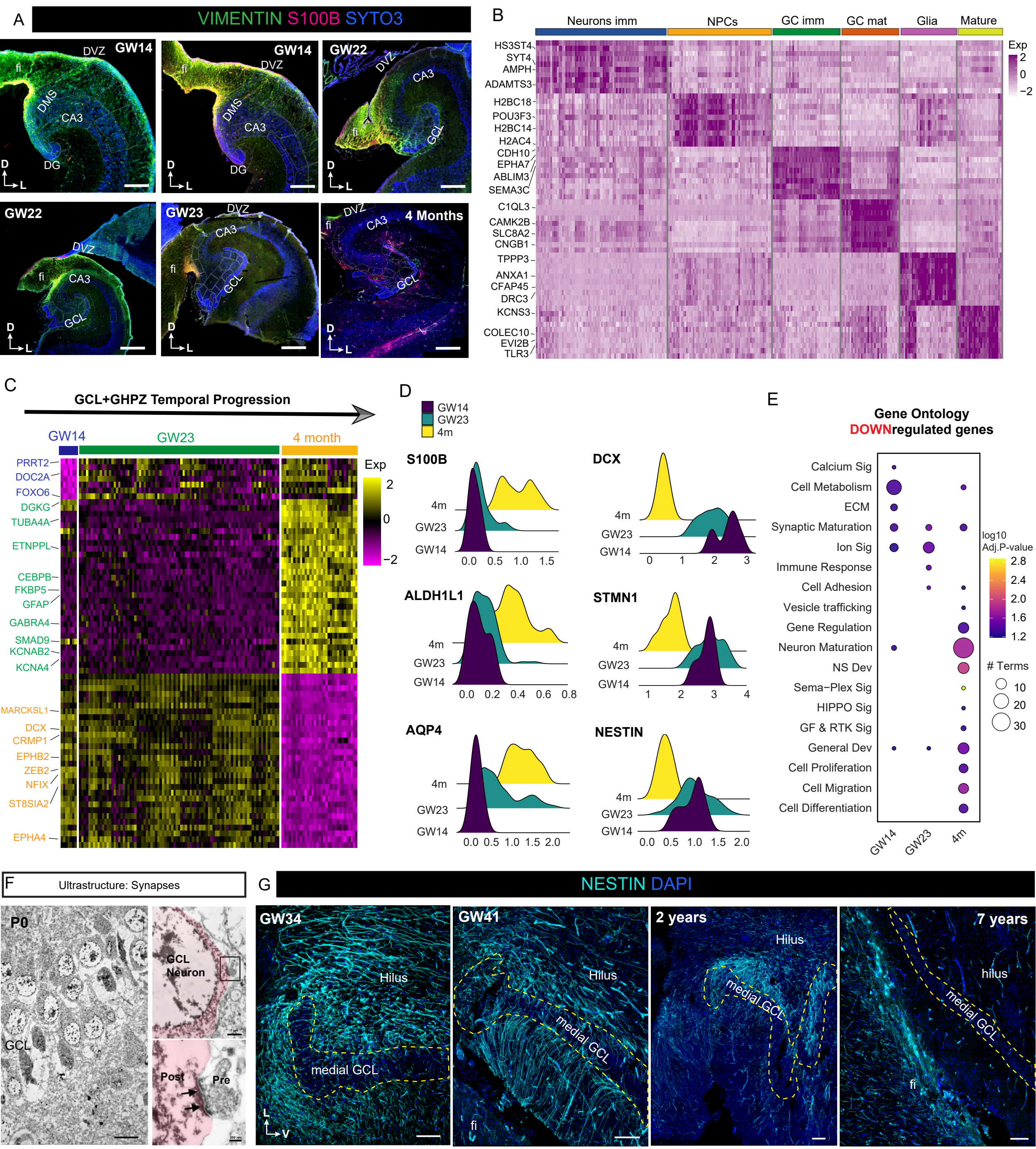

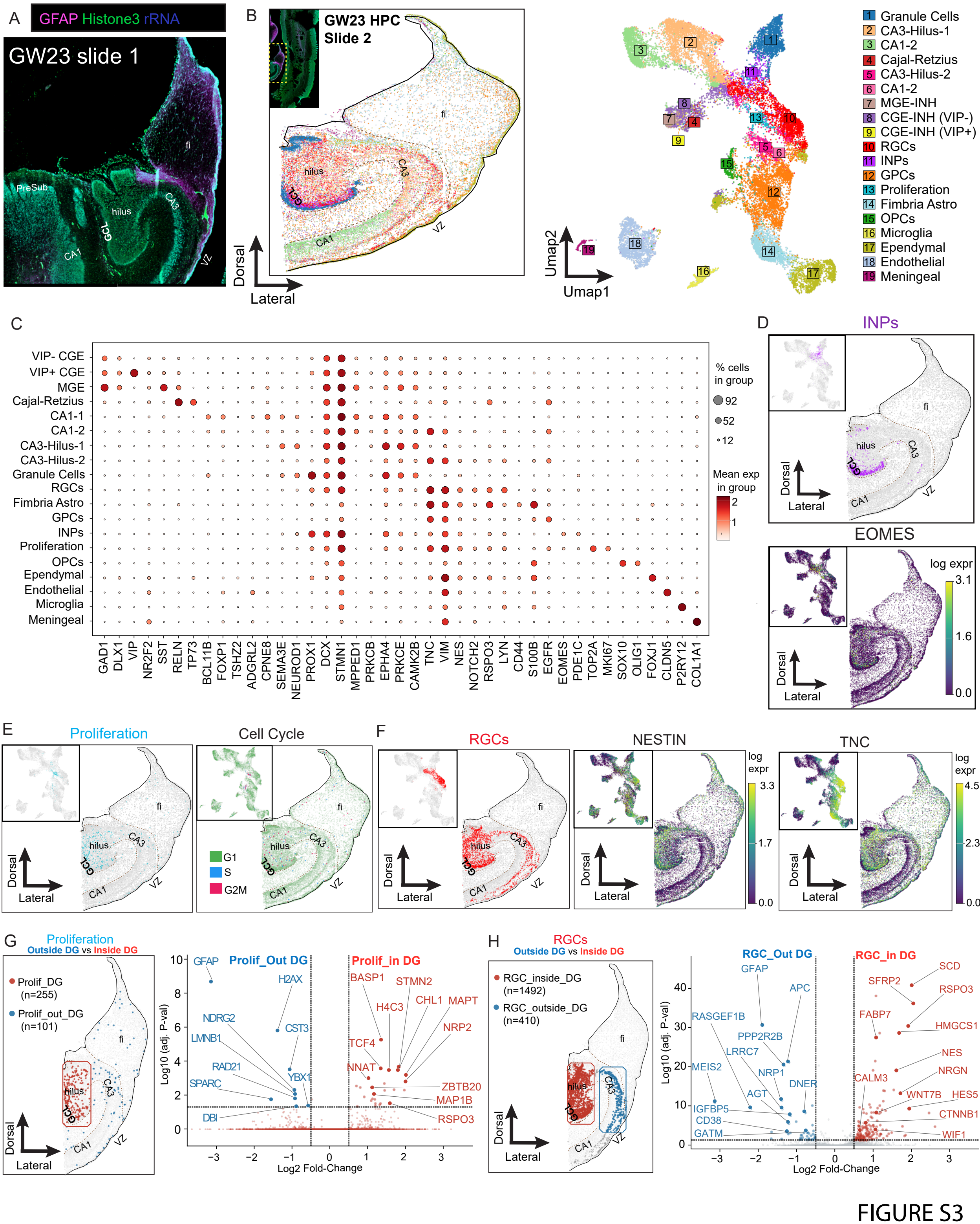

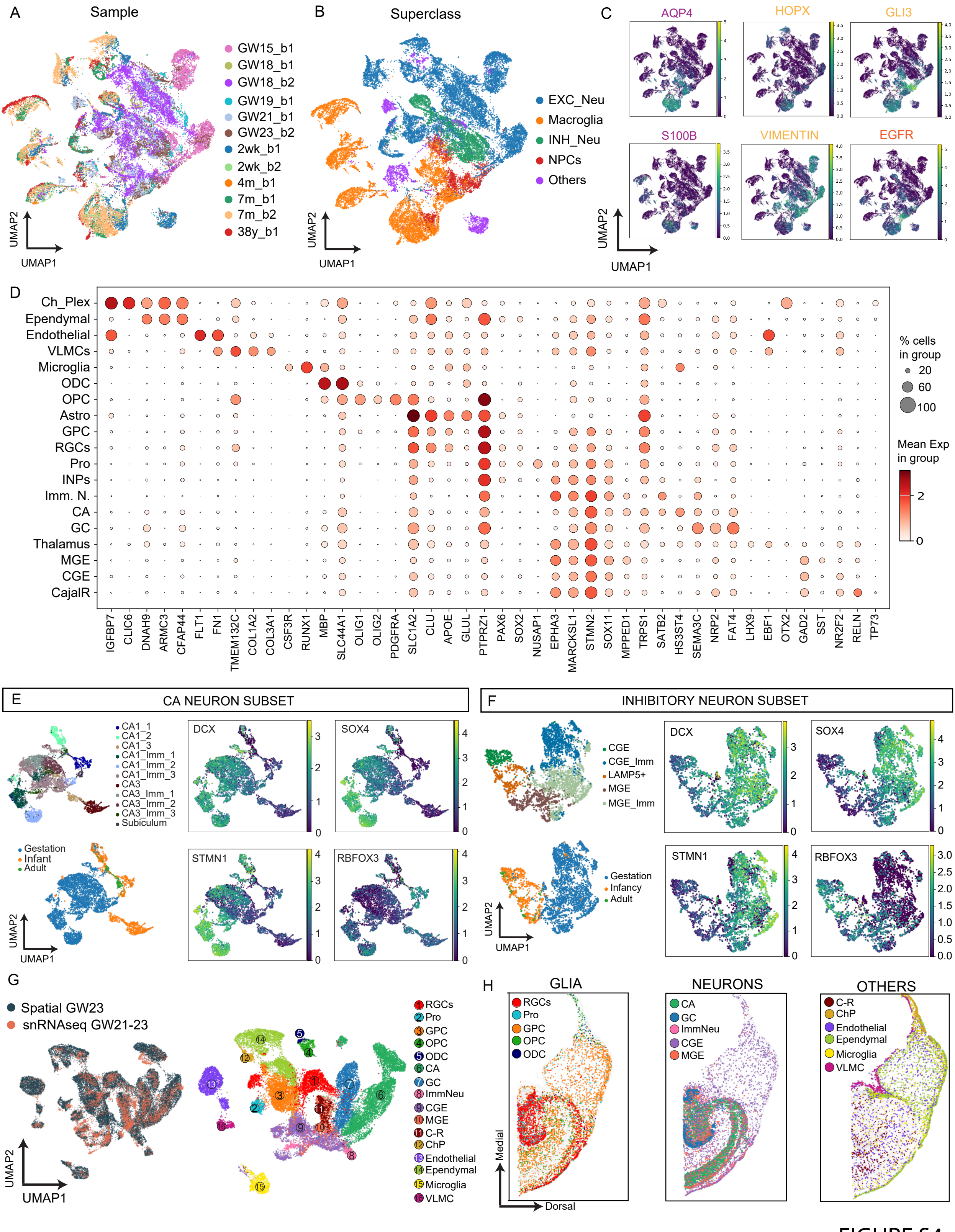

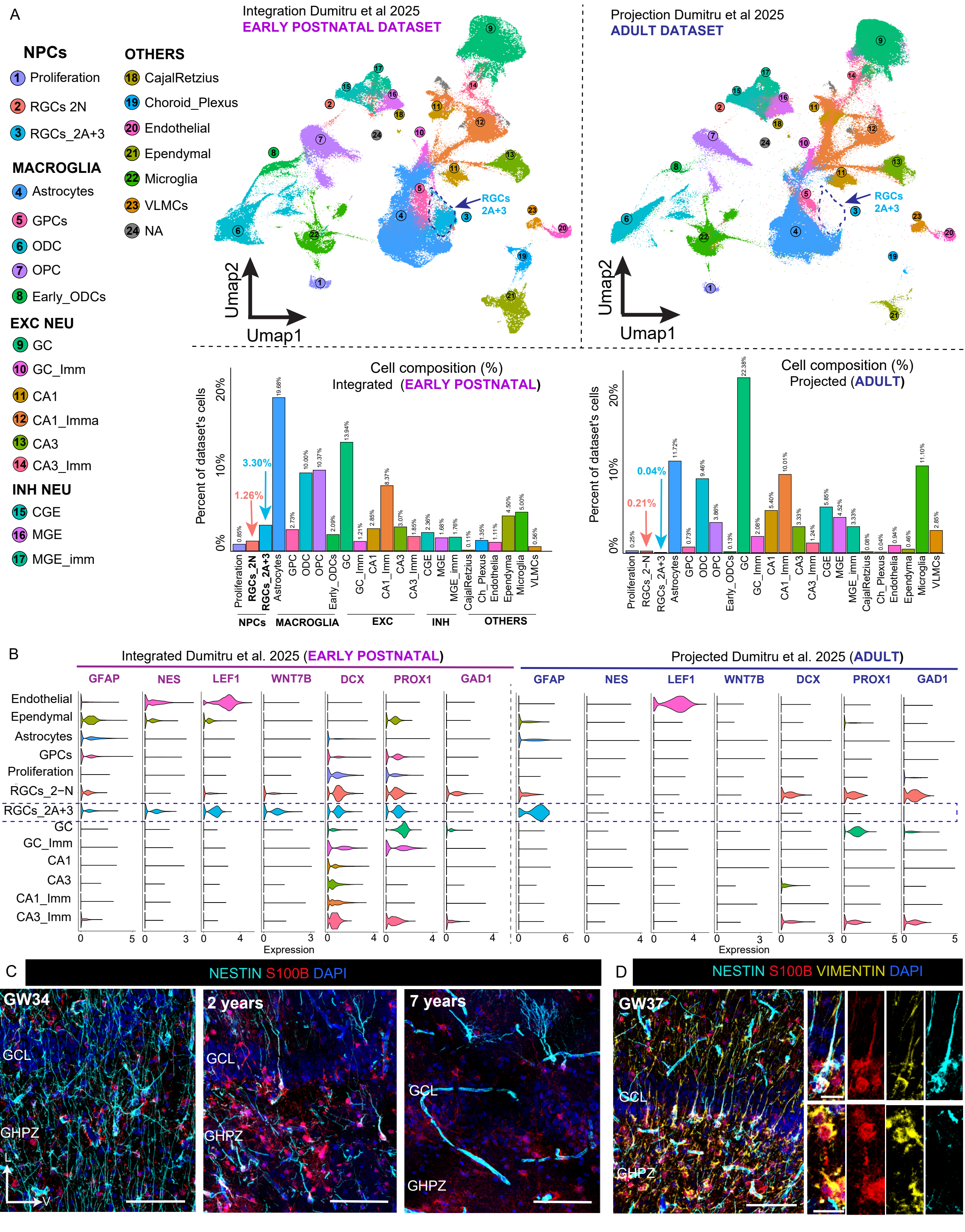

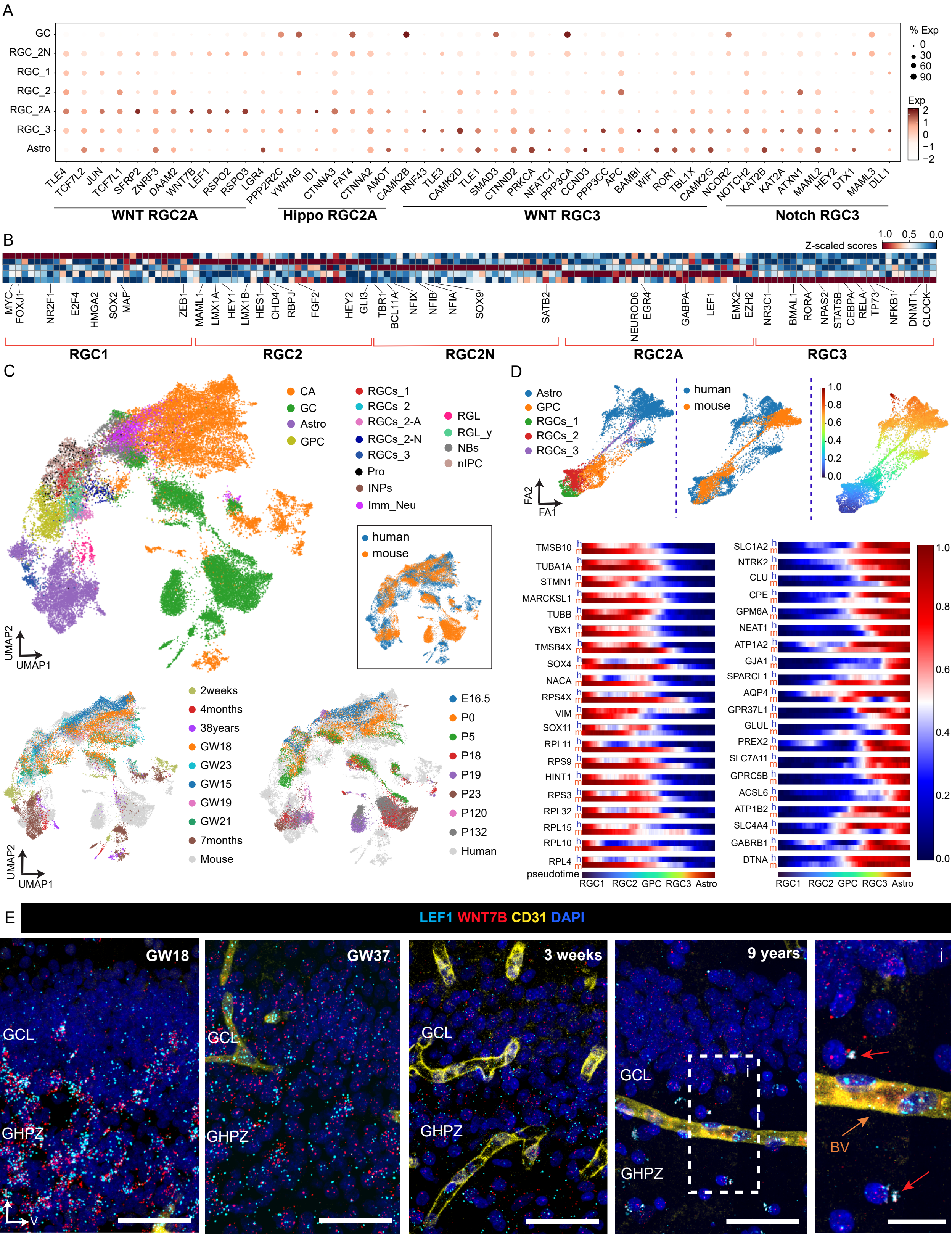
